# PepVAE3: variational autoencoder framework for antimicrobial peptide generation and activity prediction

**DOI:** 10.1101/2021.04.07.438720

**Authors:** Scott N. Dean, Jerome Anthony E. Alvarez, Dan Zabetakis, Scott A. Walper, Anthony P. Malanoski

## Abstract

New methods for antimicrobial design are critical for combating pathogenic bacteria in the post-antibiotic era. Fortunately, competition within complex communities has led to the natural evolution of antimicrobial peptide (AMP) sequences that have promising bactericidal properties. Unfortunately, the identification, characterization, and production of AMPs can prove complex and time consuming. Here we report a peptide generation framework, PepVAE3, based around variational autoencoder (VAE) and antimicrobial activity prediction models for designing novel AMPs using only sequences and experimental minimum inhibitory concentration (MIC) data as input. Sampling from distinct regions of the learned latent space allows for controllable generation of new AMP sequences with minimal input parameters. Extensive analysis of the PepVAE3-generated sequences paired with antimicrobial activity prediction models supports this modular design framework as a promising system for development of novel AMPs, demonstrating controlled production of AMPs with experimental validation of predicted antimicrobial activity.

## Introduction

Many pathogenic bacteria are resistant to the majority of, if not all, antibiotics that are currently being isolated using traditional methods. Because of this, generation of new antimicrobials is critical for survival in the post-antibiotic era (Brown and Wright 2016). At the current rate, annual global death due to antibiotic resistance is projected to exceed 10 million by 2050, costing 100 trillion USD (O’Neill 2014); however, investment has not kept up with the task with very few antibiotics currently in clinical development and far fewer likely to be approved for treatment of patients (Trusts 2019). To lower the burden required for antimicrobial development, various schemes for their discovery and refinement have been proposed, including improved methods for cultivation of antibiotic-producing organisms (Ling, Schneider et al. 2015) and repurposing of FDA-approved drugs as antimicrobials (Farha and Brown 2019). Recent work has shown that generative deep learning techniques can be applied to this problem: using a generative model, Stokes et al. showed that by training on a starting database of compounds from ZINC15 (Sterling and Irwin 2015) new small molecule antibiotics can be identified, such as halicin, which displays activity against *Acinetobacter baumannii* in a murine model of infection (Stokes, Yang et al. 2020) and is currently in clinical trials.

Unlike many small molecule antibiotics, antimicrobial peptides (AMPs), essential components of the innate immune system of humans and other organisms, have retained effectiveness as antimicrobials despite their ancient origins and widespread and continual contact with pathogens (Lazzaro, Zasloff et al. 2020). For this reason, among others, peptide antibiotics have been regularly deemed “drugs of last resort” for their ability to kill multidrug resistant bacteria, an increasingly important classification due to resistance formation towards conventional antibiotics (Lewies, Du Plessis et al. 2019). Generally acting through mechanisms associated with membrane disruption, as well as other routes of incapacitation (Chung, Dean et al. 2015), the relative immutability of bacterial membranes and other essential AMP targets make the development of resistance to AMPs rare, but possible (Kubicek-Sutherland, Lofton et al. 2016), thus increasing the importance of their reliable, continued discovery, to grow to the antimicrobial stockpile (Lazzaro, Zasloff et al. 2020).

Attempts at both generating new AMPs and improving their activity have been carried out with varying degrees of success (Mahlapuu, Bjorn et al. 2020). Many of these new or enhanced AMPs have been generated using low-throughput design methods, including rational design and specific amino acid substitution, *de novo* peptide design of alpha helices such as (LKKL)3, use of templates or motifs, or otherwise high throughput techniques such as rational library design, each of which requiring expert knowledge (Huan, Kong et al. 2020). Certain high throughput computational techniques such as genetic algorithms have shown promise; however, in many applications starting sequences are directed toward canonical amphipathic alpha-helical peptides, restricting output to a small subset of possible structures and sequences (Porto, Irazazabal et al. 2018).

In order to increase the rate of discovery of AMPs, newer high-throughput and low expertise design approaches are needed. In a similar vein to Stokes et al., several recent preprints and publications have demonstrated the application of generative deep learning on the design of AMPs, using long short-term memory (LSTM) networks (Müller, Hiss et al. 2018, Nagarajan, Nagarajan et al. 2018), Generative Adversarial Networks (GANs) (Tucs, Tran et al. 2020), and Variational Autoencoders (VAEs) (Das, Wadhawan et al. 2018, Dean and Walper 2020). These works have been facilitated by thousands of AMP sequences housed in various databases, including Antimicrobial Peptide Database 3 (Wang, Li et al. 2016) and Data Repository of Antimicrobial Peptides (Kang, Dong et al. 2019) which pair peptide sequences with experimentally determined activity against Gram-negative and Gram-positive bacteria, fungi, HIV, and cancer cells. To date several of these datasets have been formatted so the sequential amino acid residues of AMPs can be represented in the form of a string of characters enabling machine/deep learning training and analysis of these datasets to identify novel AMP sequences (Müller, Hiss et al. 2018, Nagarajan, Nagarajan et al. 2019, Witten and Witten 2019).

Although recent results using generative deep learning for producing new sequences has shown promise, including a handful of experimental demonstrations of their activity (Nagarajan, Nagarajan et al. 2018, Dean and Walper 2020, Tucs, Tran et al. 2020), some improvements are necessary. Of foremost importance, as many of these systems readily generate sequences that are both predicted and experimentally found to be inactive, machine/deep learning systems would benefit from integrated activity prediction functions. Although AMP activity prediction applications exist, many are classifiers, predicting a binary antimicrobial vs. not (Gabere and Noble 2017), where regression models would be of greater value. Additionally, a general reduction in filtering steps would reduce bias in predicted AMPs, since amphipathicity and helicity are explicitly selected for by certain models (Nagarajan, Nagarajan et al. 2018, Porto, Irazazabal et al. 2018). Thus, generation of new sequences with desired predicted activity via a semi-unsupervised and streamlined AMP-generation framework with minimal input parameters and less potential for biased output would be an improvement over previous works. In this study, we report the joining of predictive models for AMP activity (minimum inhibitory concentration (MIC)) with a generative VAE for an automated framework to produce new peptides with experimentally testable predicted activity. We demonstrate an improved automated semi-supervised approach for generating promising new sequences and experimental investigation, resulting in low MIC AMPs against *Escherichia coli*, *Staphylococcus aureus*, and *Pseudomonas aeruginosa* output from a handful of input parameters.

## Results

### Dataset characterization and framework design

This study makes use of the Witten and Witten GRAMPA dataset as the starting point (Witten and Witten 2019). Within this dataset, amino acid sequence and MIC values for peptides targeting several common bacterial species including *E. coli*, *S. aureus*, and *P. aeruginosa* are reported with *E. coli* being the most counted at 9150 different entries (**Figure S1A**). After filtering the dataset on bacteria species, peptides that were of length ≥ 40 or contain cysteine were removed in order to avoid costly and difficult synthesis of long peptides, as well as the complications cysteine-containing peptide create for their production, activity testing, and aggregation. The peptide length distribution of AMPs (without cysteine) tested against *E. coli* had a median of 17 amino acids prior to filtering on length (**Figure S1B)**, and median of 16 amino acids following removal of long sequences (**Figure S1C**). Of the remaining peptides (N = 3280), the median log μM MIC value was found to be 1.19 with a net charge of +4 (**Figure S1C-D**). Finally, to obtain a glimpse of possible secondary structures of the AMPs in this dataset, we calculated the hydrophobic moments at different angles. The noticeably higher hydrophobic moment at 100 degrees in **Figure S1E** suggests helical secondary structure likely predominates, with a relatively minor proportion of beta sheet and random coil comprising the remainder.

For the next two most common species found in the dataset following *E. coli, S. aureus* and *P. aeruginosa,* we performed the same characterization as described above. Although the overall counts were lower for the *S. aureus* and *P. aeruginosa* AMP datasets at 2974 and 1968, respectively, both the distributions and median values for length, MIC, charge, and hydrophobic moment were found to be similar to those found for *E. coli* (see **Figure S2** **and** **S3**, respectively).

Using the above defined dataset for *E. coli*, we designed a VAE AMP generation pipeline (**Figure 1**). Broadly, the VAE AMP generation and design process occurs in two stages: (1) algorithm training and (2) AMP evaluation. In the first stage, the VAE is trained on a curated AMP dataset followed by development of a regression model for activity prediction and the subsequent development of the latent space. Stage two includes the identification of new AMP sequences from the latent space (sampling) and the subsequent production and characterization of the AMPs including determination of the MIC values. A VAE implemented as previously described (Dean and Walper 2020), making use of VAE described by (Bowman, Vilnis et al. 2015), was trained on the *E.coli* dataset as described above. The number of intermediate dimensions was set to 1024 and latent dimensions was set to 50. Training was stopped after 500 epochs or when loss decreased at a sufficiently low rate. The final state of the model was saved and used for sampling novel sequences. A more detailed description of the framework design is provided in the Methods section. As demonstrated in previous work, the implicit starting assumption was that sequence order, or “peptide grammar,” and characteristics dependent on that sequence were the components “learned” by the VAE. Output of the VAE was a 50-dimensional latent space where each of the sequences is encoded to a unique location. Once generated, coordinates can be chosen from the latent space and translated to novel AMP sequences using the generated decoder (see diagram in **Figure 1**).

**Figure 1.**
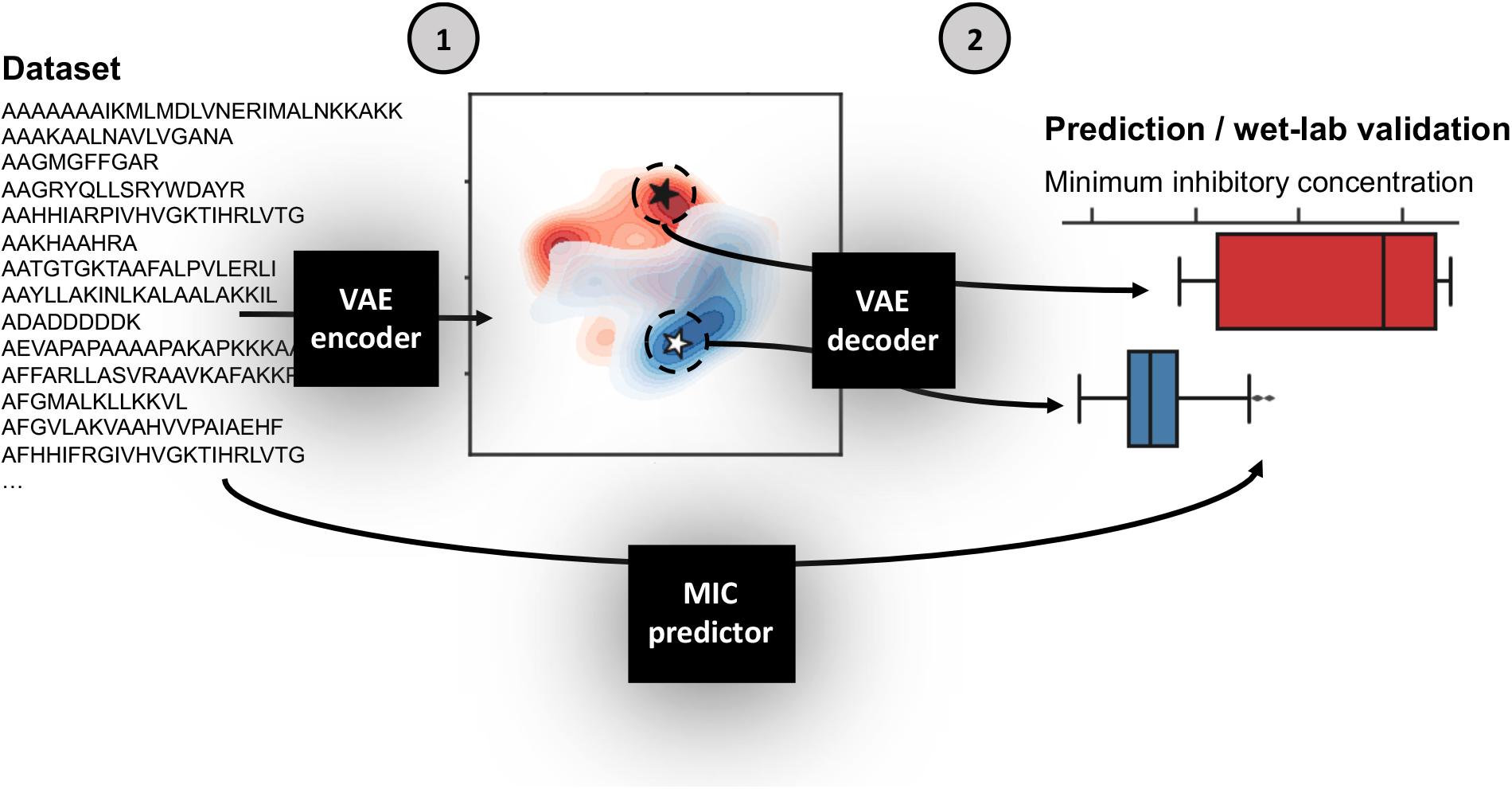
Schematic of VAE AMP generation and design process. The VAE AMP generation and design process occurs in two stages: (1) training the VAE for the development of the latent space and a regression model for activity prediction, and (2) sampling from the latent space, generation of new AMPs, and assignment of predicted MIC values. The first step of Stage 1 was to train the VAE on the *E. coli* dataset. The general design of the VAE was previously described (Dean and Walper 2020), making use of VAE described by (Bowman, Vilnis et al. 2015) which was reported for use in generating new sentences. Here, the number of intermediate dimensions was set to 1024 and latent dimensions was set to 50. Training was stopped after 500 epochs or when loss decreased at a sufficiently low rate. The final state of the model was saved; the encoder is used in Stage 1 and the decoder is used in Stage 2. The MIC prediction regression model is similarly trained on the same dataset and used in Stage 2 following sequence generation by the VAE decoder to assign MIC values for those new AMPs against *E. coli*. A more detailed description of the framework design is provided in the Methods section.

### VAE, latent space visualization, and sampling

In order to visualize the organization of the developed 50-dimension latent space, multiple dimensionality reduction techniques were tested: principal component analysis (PCA), T-distributed stochastic neighbor embedding (t-SNE), and Uniform Manifold Approximation and Projection (UMAP) (**Figure S4A**). Upon visual inspection, t-SNE and UMAP show separation between the distant MIC thresholds of < 0.2 log μM and >2 log μM, while separation between the two groups in the first two components of the PCA is less clear; this is supported using Adjusted Rand Index (ARI) measurement and Adjusted Mutual Information (AMI) scores (**Figure S4B**). Here, 2D projects (via PCA, t-SNE, and UMAP) of the latent representation was used as input to the *K*-means algorithm and measure the overlap between the resulting clustering annotations and the pre-specified subpopulations (the < 0.2 log μM and >2 log μM labels) using the Rand index and AMI scores. By ARI, t-SNE shows the highest separation measure with 0.62, with UMAP at 0.58, and PCA at 0.47. Using AMI score, t-SNE is also highest at 0.59, t-SNE at 0.56, and PCA at 0.47. These results suggests that 1) the AMPs encoded to the latent space are not randomly distributed in terms of their MIC value classification, and 2) t-SNE provides superior 2D clustering visualization in this application, relative to PCA and to a lesser extent UMAP. t-SNE projections are shown in **Figure 2A-B.**

**Figure 2.**
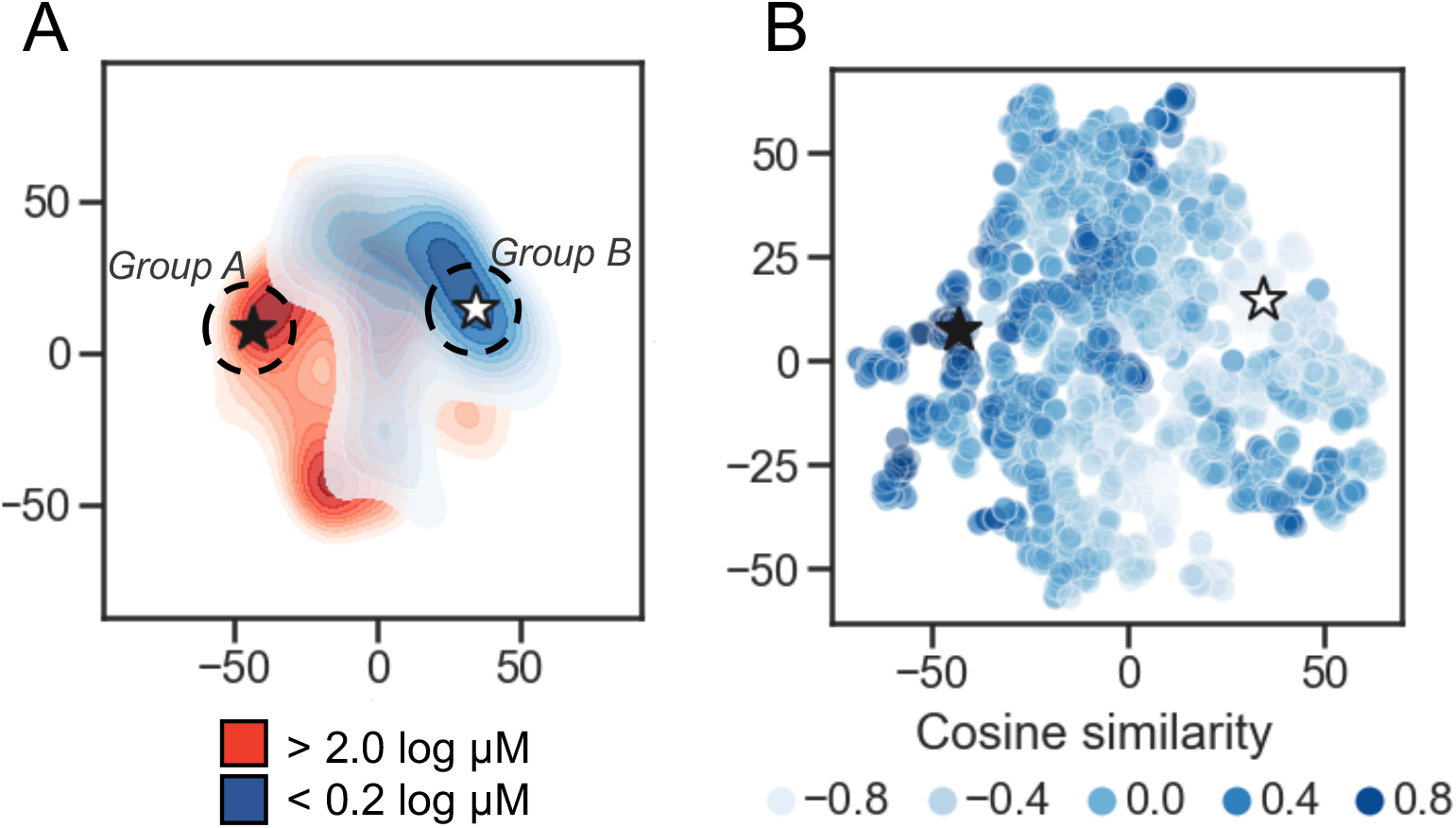
Latent space characterization. Dimensionality reduction for visualization of the 50-dimension latent space. A) 2D contour plot of t-distributed stochastic neighbor embedding (t-SNE) with two components performed on the encoded peptides. The MIC thresholds for coloring were: < 0.2 log μM is shown in blue, >2 log μM shown in red, and those with values ≥ 0.2 and ≤ 2 were set to light gray. Regions of higher density are darker. The black star is located at the peptide caseicin B-B1, and a white star is located at the embedding most distant in cosine similarity (encoding for the peptide trialysin peptide P4). Group A and Group B sampling locations are schematically shown at the dashed circle around caseicin B-B1 and trialysin peptide P4, respectively. B) Scatterplot of t-SNE projection showing encoded peptides colored by cosine similiartity calculated using the vector encoding for peptide caseicin B-B1 (black star) as reference. Higher similarity values indicate more similarity between vectors; lower values indicate more difference.

To delve into the organization of specific AMPs encoded to the latent space, we used cosine similarity as a measure of distance between AMPs using their 50-dimension vectors as input. First, the cosine similarity for all vectors was calculated relative to each vector, generating a similarity list for every AMP. For each similarity list, the associated MICs for the five most similar vectors were averaged; this process was repeated for the five least similar vectors, and the difference between the two averages was taken. From this process, the greatest difference highlighted a largely inactive AMP, VLNENLLA, called caseicin B variant B1, a variant of caseicin B which is found in milk. Regardless of mutation, caseicin B exhibited a MIC of approximately 1250 μM against *E. coli* NCIMB 11843 (Norberg, O’Connor et al. 2011). Since this variant of caseicin B and other mutants each showed low activity against *E. coli* (≥ 1.25 mM), we were confident in sampling from latent space near their location would likely produce new AMPs similarly inactive and hypothesized that AMPs generated in a region most distant from this reference point were likely to be highly active against *E. coli.* Caseicin B-B1 is identified in **Figure 2A-B** at the black star; a white star is located at the embedding most distant in cosine similarity encoding for the peptide KFGKIVGKVLKQLKKVSAVAKVAMKKG, trialysin peptide P4. Trialysin peptide P4 is a potent pore-forming peptide found in the saliva of *Triatoma infestans*, the insect vector of Chagas’ disease, and lytic to *E. coli* in LB broth at approximately 10 μM (Martins, Sforça et al. 2006). In **Figure 2A**, both caseicin B-B1 and trialysin peptide P4 locate within regions of low activity and high activity AMPs as organized within latent space, respectively, and in **Figure 2B** both encode to regions of similarly high and low cosine similarity.

Next, we took the highest and lowest five AMPs (by cosine similarity) and grouped them: parent Group A and parent Group B. Here, the parent Group A five AMPs (closest to and including caseicin B-B1) were identified as VLNENLLA, VLNENLAA, VLNENLLK, VLNENLL, and VLNENLLH, each of which are caseicin B variants reported by Norberg *et al.* (Norberg, O’Connor et al. 2011). And parent Group B (most dissimilar to caseicin B-B1) was: KWKLWKKIEKWGQGIGAVLKWLTTWL, KWKSFIKKLTSAAKKVVTTAKPLISS, KWKSFIKKLTSVLKKVVTTAKPLISS, KFFKKLKKAVKKGFKKFAKV, and KFGKIVGKVLKQLKKVSAVAKVAMKKG. The average MIC against *E. coli* for parent Group A group was 2500 μM, and for parent Group B: 1 μM. Nearby these ten AMPs (two groups of five) a total of 100 peptides were generated by the decoder. Following removal of duplicates or those that are already present in the database, 38 remained and were synthesized (see sequences in **Table 1**), with 22 peptides in the Group A and the remaining 16 peptides were in the Group B. The 38 sequences were designated names p1-38 and were associated with Parent peptide IDs corresponding to those Peptide IDs found in **Table S1** – the dataset used for training the model. Cosine similarity listed in **Table 1** is relative to caseicin B variant B1. For purposes of comparison, in addition to the 38 peptides from Group A and Group B, 100 control sequences were decoded from random 50-dimension vectors.

**Table 1.**
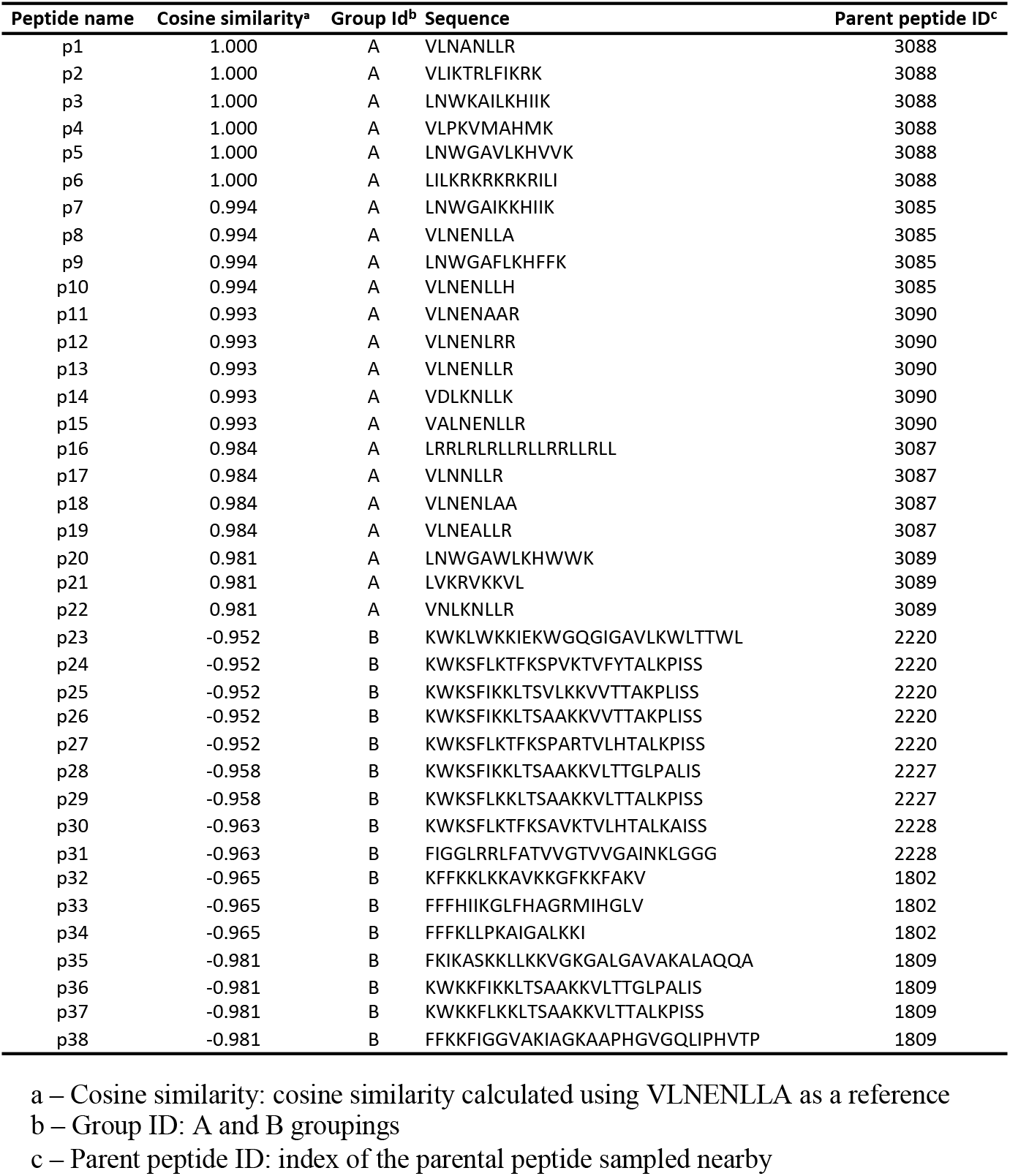
PepVAE3-generated peptides.

### AMP generation, characterization, and MIC results

A secondary structure prediction algorithm (GOR IV) was used to predict helix, sheet, and coil percentages of the Group A and Group B sampling groups. Group A peptides were predicted to have similar proportions sheet and coil with medians 30% sheet and 37% coil, with a median of 0% helix (**Figure 3A**). Conversely, Group B peptides were predominately helical at 62%, with the remainder composed of approximately equal proportion sheet and coil. Group A and Group B peptides are significantly different for both predicted proportion of helix and coil (p < 0.01, Welch’s two-sided t-test). For comparison, the group of 100 random sequences were not found to be statistically different from Group A, while Group B had significantly higher in percent helix and significantly lower in percent coil (p < 0.01, Welch’s two-sided t-test) (**Figure S5A**). Given the relatively high proportion of peptides predicted to be helical in Group B, the amphipathic nature of both groups was examined via calculated hydrophobic moments at 100 degrees. Predictably, differences between Group A and Group B are significant (p < 0.01, Welch’s two-sided t-test). As expected, the distribution of hydrophobic moment of the randomly generated group was similar to that of the sequences found in the training set when the hydrophobic moment at 100 degrees is calculated, suggesting the VAE generations aligned well with real data (see **Figure S1F** and **Figure S5B**). Altogether these results suggest the 22 Group A peptides are predicted to be significantly less helical than the 16 Group B peptides, while randomly sampling captures a wider range of predicted structures and non-amphipathic/amphipathic AMPs. Importantly, the predictions suggest controlled sampling from distinct subpopulations of latent space generates sequences with significantly different characteristics.

**Figure 3.**
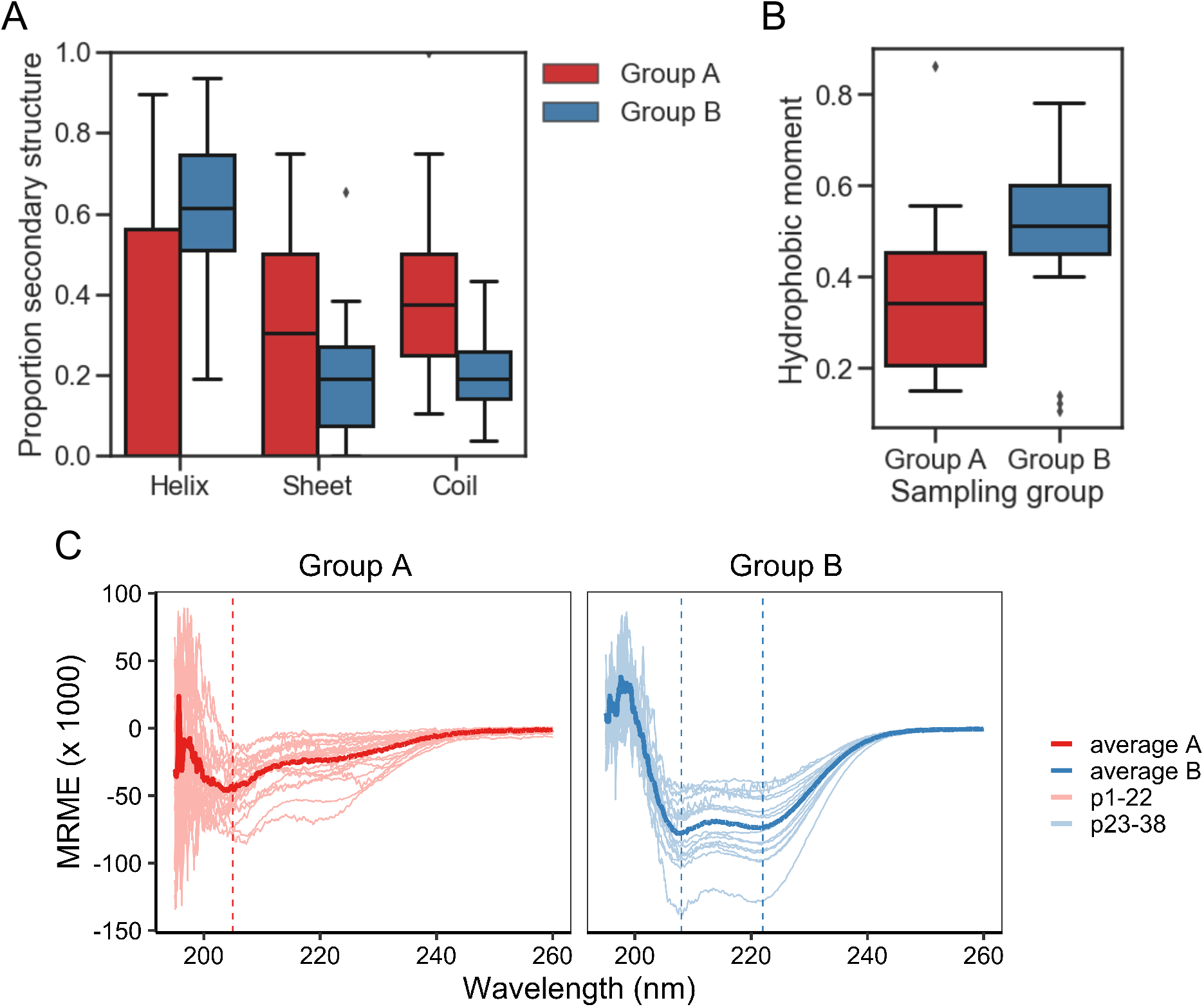
Generated peptide characterization. A) Boxplot of the predicted helix, sheet, and coil percentages calculated from Group A and Group B sampling groups into the GOR IV algorithm. Group A and Group B are significantly different (p < 0.01, Welch’s two-sided t-test) for helix, sheet, and coil. B) Boxplot of the calculated hydrophobic moments obtained by applying the modlAMP function calculate_moment on the sequences from Group A and Group B sampling groups. The differences between Group A and Group B groups are significant (p < 0.01, Welch’s two-sided t-test). C) Mean residue molar ellipticity (MRME) plots of peptides from group A (left) and group B (right) in the presence of membrane mimic 60 mM SDS. Each scan was averaged from three scans for each peptide with peptide-free buffer baseline scan subtracted. Lighter colored lines are individual peptide scans; darker colored lines are average scan for the group. Vertical lines highlight approximate minima of average scans: 205 nm for Group A; 208 and 222 nm for Group B.

Experimental investigation of secondary structure was performed using circular dichroism (CD) in phosphate buffer with sodium dodecyl sulfate (SDS) micelles as a membrane-mimicking agent (Tulumello and Deber 2009). To account for concentration and difference in peptide length, mean residue molar ellipticity (MRME) was plotted to visualize the relative proportion of secondary structure for each group A and B (**Figure 3C).** Results in the presence of SDS show that the average Group A peptide presents a mixture of random coil and helical character with minima predominant at ~205 nm suggests a largely random structure, with a smaller but noticeable dip at 222 nm suggesting a minor percentage of helix. Individual scans separated out (**Figure S6A**) shows a mixture of structures. Group B peptides were predominantly helical, with paired minima at ~208 and ~222 nm and, unlike Group A, were more uniform in the scans of individual peptides. Using the circular dichroism analysis program Beta Structure Selection (http://bestsel.elte.hu/), secondary structure was estimated from the CD data (converted to Δ*ε*). Following summation of antiparallel, parallel, and turn into “sheet”, the results were plotted for both Groups A and B (**Figure S6B**). Analysis showed that Group A and B with median percent helicity of 4% and 63%, respectively. The results are comparable to those estimated from sequence via GOR IV.

Although, secondary structure and particular measurements such as hydrophobic moment are closely related to the antimicrobial activity of AMPs, particularly those with alpha helical character, more predictive measures are available in the form of regression models. Witten et al. and others have reported use of regression models, including a convolution neural network (CNN) (Witten and Witten 2019). In our study, we implemented the reported CNN as well as several machine learning regression models for MIC prediction against *E. coli.* Preliminary tests utilized model frameworks within the Statistics and Machine Learning Toolbox and Regression Learner app in MATLAB (**Figure S6**). However, the long training time of the best performing model identified (Rational Quadratic Gaussian Process Regression) led us to migrate to other models implemented in Python. The CNN from Keras, elastic net, gradient boosting (GB), kernel ridge, lasso, and random forest (RF), each from Scikit-learn, as well as light gradient boosting machine (LGBM) and an extreme gradient boosting (XGB) model were initially tested. Our results showed that three of the examined models significantly underperformed the others: lasso, kernel ridge, and elastic net, each with R^2^ < 0.4 and relatively high RMSEs (**Figure S8A**). This underperformance was also visible in the actual-predicted difference histograms (**Figure S8B**) where each of their distributions were flatter than the others. For this reason, these three models were excluded from further study. In addition, the relatively complex CNN implemented as described by Witten *et al.* in Keras was underperforming in relationship to the length of time required to train the model and was therefore also excluded. The remaining four models – GB, RF, LGBM, and XGB – were further interrogated. Representative scatterplots are shown in **Figure 4A**, with each showing R^2^ higher than 0.67. Following successive train-test split-shuffling cross validation, GB was found to have both the highest median R^2^ (0.73) and lowest RMSE (0.50) (**Figure 4B-4C**) and was used going forward for MIC prediction of AMPs against *E. coli.* Likewise, using the *S. aureus* and *P. aeruginosa* datasets, we identified the best-performing models for predicting MIC against both *S. aureus* (**Figure S9**) and *P. aeruginosa* (**Figure S10**). For *S. aureus* XGB was found to be the best predictor after cross validation (**Figure S9**), while for *P. aeruginosa,* although the RSME and R^2^ measures disagreed, the RF model was selected (**Figure S10**). Although several regression models for MIC prediction of AMPs on *E. coli* have been reported with varying degrees of accuracy (Wu, Wang et al. 2014, Xiao and You 2015, Nagarajan, Nagarajan et al. 2018, Gull 2020), none have publicly accessible models in order to directly compare with our results. Nevertheless, given the modular nature of the described PepVAE3 framework, any superior MIC prediction system could be used in place of the described models.

**Figure 4.**
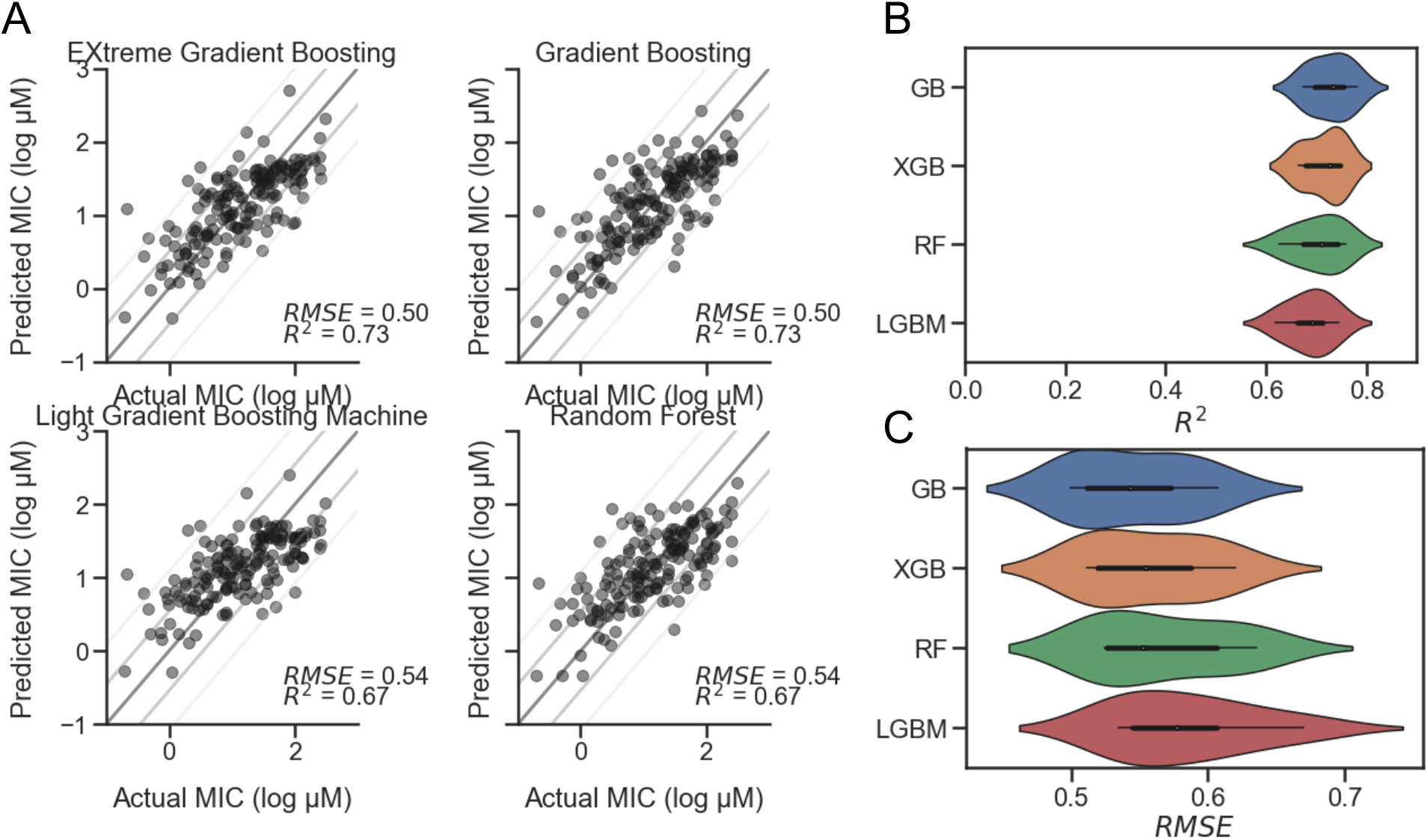
Comparison of MIC prediction models. Four regression models for predicting AMP MIC values against *E. coli*. A) Representative scatterplots of Predicted vs. Actual MIC (log μM) of EXtreme Gradient Boosting (XGB), Gradient Boosting (GB), Light Gradient Boosting Machine (LGBM), and Random Forest (RF) predictions on holdout test dataset with RMSE and R^2^ values displayed. Lines represent standard diagonals, in addition to the diagonal +/− 1 and 2 standard deviations of the points. B) Results of cross validation using shuffled split (n=25) shown as a violin plot for each model, sorted from highest to lowest mean R^2^ value, with GB at the top with a median of 0.73. C) Cross validation results for RMSE sorted from lowest to highest mean RMSE value, with GB at the top with a median of 0.54.

MICs predicted for Group A and Group B peptides are provided in **Figure 5** and **Table 2**. For *E. coli*, the median predicted MIC of the Group A group was 1809 μM and 2 μM for Group B, and the sets were found to be significantly different (p = 1 × 10^−8^, Welch’s two-sided t-test). As expected, the median of predicted MICs for the peptides decoded from randomly selected points in latent space was in between groups A and B: 12 μM (**Figure S5C**). These predicted results are similar to the MICs of the parent AMPs used for generation of Group A and B. To investigate the predicted MICs of an intermediate location, we filtered regions of the latent space by cosine similarity relative the caseicin B variant B1 reference, randomly sampled 10 sequences, then generated new sequences (n=10) around these using the same method as described above and predicted the MICs for each group (**Figure S11)**. Each group was found to be significantly different from the other (p < 0.01). These results suggest intermediates locations between the polar ends of cosine similarity (and between Group A and Group B) would likely on average have corresponding intermediate MICs.

**Table 2.**
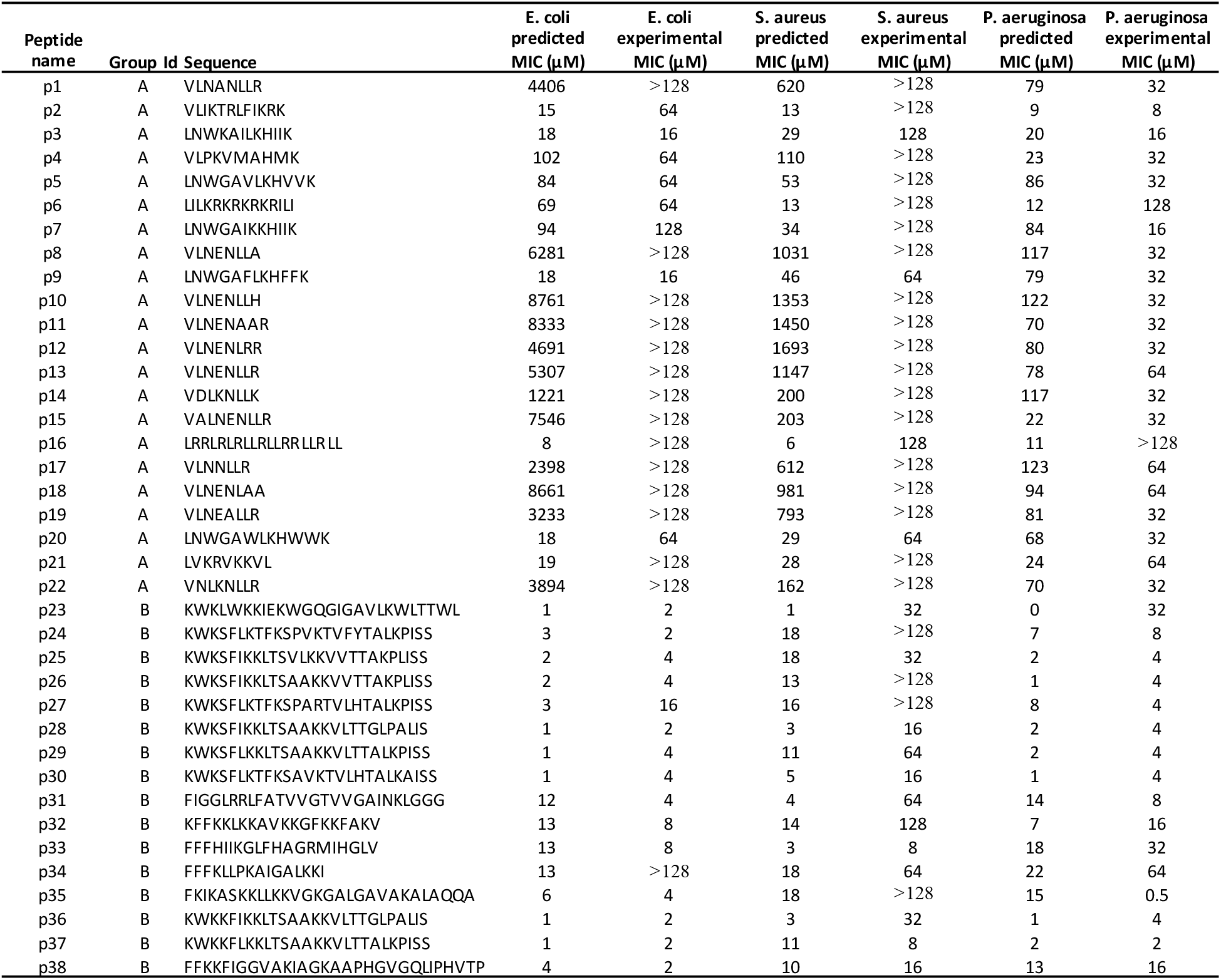
MIC assay results.

**Figure 5.**
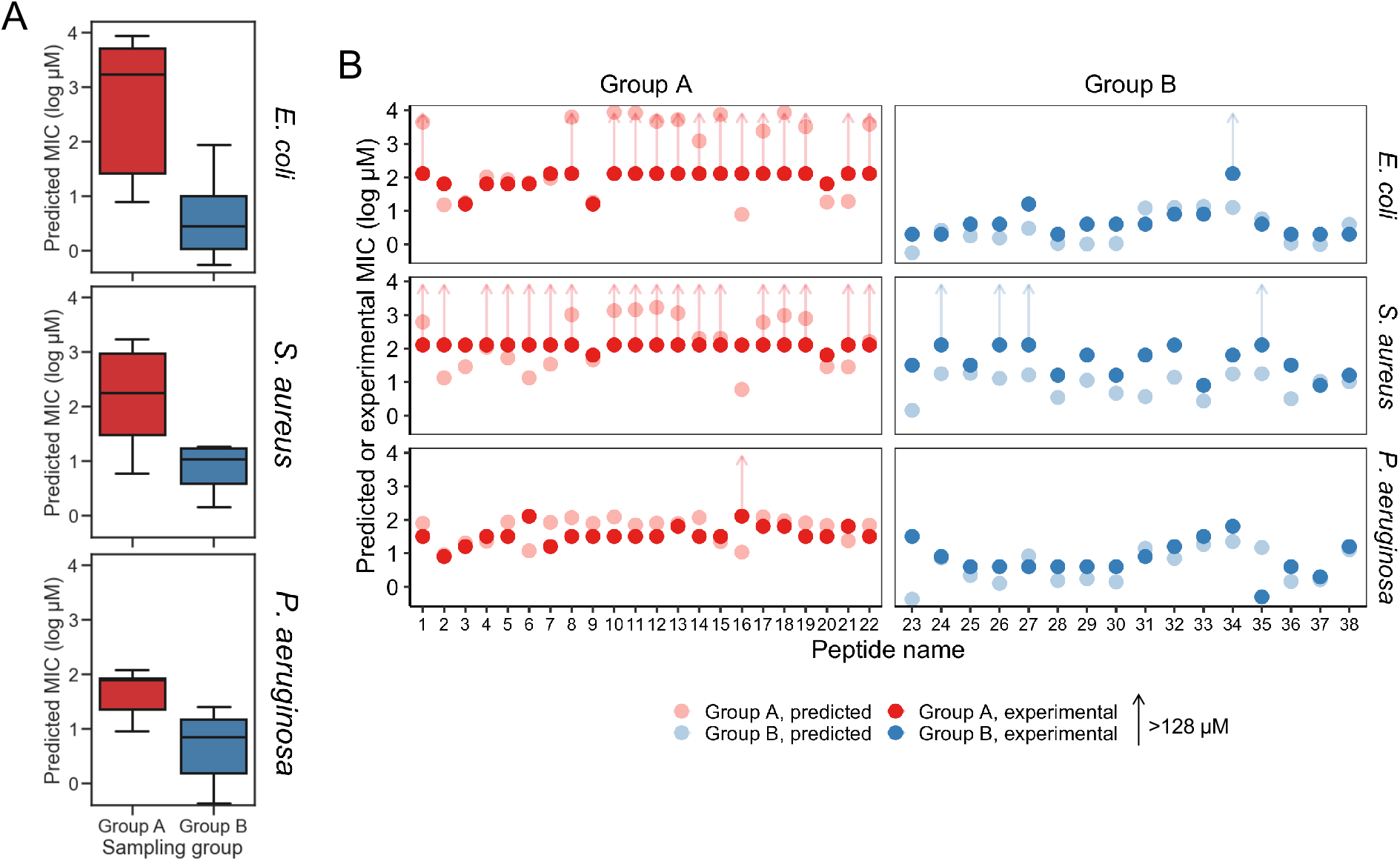
Predicted and experimental MICs against *E. coli*, *S. aureus,* and *P. aeruginosa*. A) Predicted MICs using the top-performing models for the Group A and Group B sampling groups are show in a boxplot, for *E. coli*, *S. aureus*, and *P. aeruginosa*, respectively. Group A and Group B are significantly different for each of the species (p < 0.01). B) Experimental MICs of Group A and Group B peptides against *E. coli*, *S. aureus*, and *P. aeruginosa*, with predicted MICs shown (with lighter coloring) for reference. Experimental MICs resulting in >128 μM determinations are shown with arrows extending up from 128 μM. All values are provided in Table 2.

While the VAE was trained on the *E. coli* dataset, and sampling was performed with activity against *E. coli* in mind, we additionally examined the effectiveness of the generated AMPs on *S. aureus* and *P. aeruginosa,* the two most common bacteria in the modified GRAMPA dataset after *E. coli* (**Figure S1A**). For *S. aureus*, the median predicted of Group A was 181 μM and 11 μM for Group B (**Figure 5**). A Welch’s two-sided t-test on Group A and B found p = 1 × 10^−1^. Meanwhile against *P. aeruginosa* the median predicted of Group A was 78 μM and 5 μM for Group B (**Figure 5**). A Welch’s two-sided t-test indicated a significant difference with p = 3 × 10^−7^. For both *S. aureus* and *P. aeruginosa,* random sampling yield median predicted MICs of 13 μM and 14 μM, respectively (**Figure S5C**).

To experimentally investigate the MIC of each of the 38 synthesized peptides against *E. coli*, the peptides were diluted in Mueller Hinton broth with a constant number of bacteria. Following incubation, we found that consistent with above predictions, the recorded MICs between the two sampling groups were significantly different. Among Group A AMPs, 63% of the MICs were found to be greater than 128 μM, while none of the 16 Group B AMPs were found to have MICs above 16 μM, other than peptide p34 (**Table 2**). After sorting for activity, the median value for Group A and Group B, was found to be >128 μM and 4 μM, respectively, which may be in line with the median predicted values of 1809 μM and 2 μM, although determining the accuracy of the values for Group A that are over 128 μM was not possible due to solubility issues. Importantly, the MIC predictions were not found to be different or independent from the experimental MICs, when categorized into >128 and ≤128 μM (p < 0.01, Fisher’s exact test).

The MIC results paired with secondary structure estimates from circular dichroism experiments, highlight a number of active AMPs within Group B were composed of a low (< 50%) proportion of helix, including peptides p23, p24, p27, and p38. In addition, we found that both generated AMPs p31 and p33 have net charge ≤ 3, which in similar generative studies including Nagarajan et al and others would have been placed below their threshold for proceeding to experimental testing of activity (Nagarajan, Nagarajan et al. 2018). Similarly, within Group A, there was a low-activity peptide with high helicity (> 50%) for p16, and relatively low-activity peptides with high net charge.

The experimentally determined MICs against *S. aureus* and *P. aeruginosa* were – unlike *E. coli* – further from their respective MIC predictions. For *S. aureus*, while 12 AMPs in Group A were predicted to have MICs >128 μM, 18 of the 22 peptides were found to have MICs at that level experimentally. The median MICs for Group A and Group B was found to be >128 μM and 32 μM, respectively, while the median predicted values were 181 μM and 11 μM. Similar to *E. coli*, the MIC predictions for *S. aureus* were similar, consistent with experimentally determined MICs when categorized into >128 and ≤128 μM (p < 0.01, Fisher’s exact test). Although more of the predictions and experimental results were not in agreement, an insignificant difference was found using receiver. Receiver operating characteristic analysis was performed to evaluate for classification predictions for both species, which resulted in area under the curve values for *E. coli* and *S. aureus* at 0.93 and 0.9, respectively (**Figure S12**). For *P. aeruginosa*, the median MICs for Group A and Group B were experimentally determined to be 32 μM and 4 μM, respectively, compared to median predicted values of 78 μM and 5 μM. Consistent with the predicted MICs, the experimentally determined values too showed the larger separation between the median MICs for Groups A and B for *E. coli* than for *S. aureus* and *P. aeruginosa* (**Figure 5A**; **Table 2**). When predicted and experimental MIC determinations found in **Table 2** are plotted relative to one another (**Figure 5B**), after accounting for the >128 μM inequalities the predictions and experimental results closely align for *E. coli* and *P. aeruginosa*, with the exception of a few peptides deviating beyond a 5-fold difference in experiment vs. prediction, notably p34 tested against *E. coli* highlighted above. Peptide 16 for both *E. coli* and *P. aeruginosa* was also found be inactive while having a low predicted MIC. In contrast, for *S. aureus,* experiments notably deviate from predicted MICs for many of the peptides, where most of Group B predicted MICs significantly overestimated activity.

## Discussion

This study reports the use of a peptide generation framework, PepVAE3, for discovery/design of new AMP sequences. Using the learned latent space, we demonstrate the identification of new active AMPs using reference peptides as input. Paired with antimicrobial activity prediction models, this modular framework shows the ability to produce AMPs with both predicted and experimentally validated activity against the targeted bacteria. Previous work on generative deep learning models for the design of AMPs has demonstrated the ability of VAEs to form a well-organized, usable latent space representation from which novel peptide sequences can be generated (Das, Wadhawan et al. 2018, Dean and Walper 2020). These reports when paired with similar promising work on the discovery of small molecules using VAEs and other generative deep learning models broadly establishes these techniques as valuable new methods for molecular and material design (Sanchez-Lengeling and Aspuru-Guzik 2018). Although generative deep learning models for AMP generation have previously shown their ability to produce distributions of characteristics that closely match the databases of sequences on which they are trained, unconditional generation alone does not readily solve the problem of discovering new AMPs that are potent against target bacteria. In particular, previously described systems would benefit from certain functions to improve automated discovery of new potent AMPs: testable activity prediction, a mechanism for AMP generation via a reference peptide for controlled sequence generation, and an expanded sequence space from which to sample.

Many previously described systems readily generate sequences that are both predicted and experimentally determined to be inactive, by nature of the models used when training sets include inactive or low-activity sequences. In the literature, while classifiers for AMP activity are widely available and meta-analyses of their performance has been reported (Gabere and Noble 2017), regression models for activity prediction – MIC, half maximal effective concentration, or other metrics – are less common. Several studies make use of the CAMPR3 predictor (Waghu, Barai et al. 2016) or other similar models that output abstract activity predictions as a probability (*probability(active AMP)*), not readily relatable to MIC or another antimicrobial activity metrics (Müller, Hiss et al. 2018, Nagarajan, Nagarajan et al. 2018, Dean and Walper 2020). Beyond the lack of testability of these predictions, it likely that the increasing hydrophobic moment yielded higher probabilities in the CAMPR3 predictor and other models suggesting that simply alternating groups of positively ionizable and hydrophobic amino acids will score highly, highlighting the importance of experimentally verifying the antimicrobial activity of generated AMPs. To address these issues, our study implemented ML models trained on sequence data and adjoining experimental MIC values to predict MICs of new peptide sequences. Critically, we found that the MIC predictions for *E. coli* were not statistically different from the experimental MICs and can, therefore, be used for assessing whether a potential AMP is likely to be active against *E. coli* prior to peptide synthesis. While the MIC predictions and for *S. aureus* and *P. aeruginosa* were similarly found to be statistically similar to the experimental outcomes, the inaccuracy of the predictions for MICs against these bacteria relative to *E. coli* was not unexpected due to the smaller training sets and lower accuracy of the MIC predictions on the test set. Increases in both data quantity and quality, with respect to the specific strain used, may significantly improve MIC predictions, especially for non-*E. coli* bacteria. Pairing with PepVAE3, further compilation of data for would likely allow for improved generation of AMPs targeting species of interest.

Generative models for AMP design can be bifurcated into those that use a starting sequence and those that do not (i.e., *de novo* design). To address the development of a mechanism for selecting a reference AMP for controlled sequence output, as opposed to unconditional generation, which would improve existing AMP generation frameworks, we implemented a simple AMP generation by reference method that uses limited input parameters: reference peptide selection and the number of new sequences to generate. We show that this method produces peptides with similar MICs as the input reference peptides, but with novel sequences not found in the training set. One possible limitation of this system is the likely requirement of confidence in sampling in the region arrived at by density of nearby, similar AMPs with similar activity. In our testing, we generated AMPs in reference to caseicin B-B, encoded nearby a series of single and double mutants with similar activity. It is possible that an isolated, single reference peptide is not sufficient, and that a small panel of AMPs may be required for confidence – sampling from a sparsely populated region will likely produce results similar to that of the random sample group in this study, who’s predicted MICs span a wide range.

Another issue our study improved upon is the expanded sequence space from which to sample. Other groups have shown the benefits of tying a continuous activity prediction to output from an AMP-generating models rather than simple classification, following up with experimental validation. These methods, however, used predicted amphipathicity, MIC, and charge filtering steps in order to obtain target active sequences (Nagarajan, Nagarajan et al. 2018, Porto, Irazazabal et al. 2018), restricting sequence output to a specific set of positively charged, alpha helical peptides. Our results showing that our list of newly generated active peptides includes non-canonical AMPs of low helicity and low net charge supports using the described VAE method, without imposing thresholds on peptide characteristics or otherwise biasing output post-sequence generation. However, we did perform pre-training biasing due to cost and synthesis limitations on both length and cysteine presence, and we expect the structural diversity of generated AMPs to have been greater had these limitations not been a factor, and we plan to avoid more of these biasing factors in future studies.

In addition to investigating AMPs generated using models trained on data without length or cysteine constraints, given sufficient funds for synthesis costs, future studies will examine our results that suggest sampling from nearer regions (i.e., not highest or lowest code similarity but interpolations in between) would on average result in generated AMPs with middling activity in a range between the polar Groups A and B. This interpolation may look similar in outcome to our proof-of-concept report on *de novo* generation of short ≤ 12-mer peptides using a VAE (Dean and Walper 2020) but would further validate that the VAE is producing a smooth, well-organized latent space. Another interesting feature to investigate in the future, relating to the ability to sample nearby a particular selected AMP, are the characteristics best retained by the newly generated AMPs. The results here suggest the average MIC of the generated AMPs from both Group A and Group B is similar to that of those AMP encoded near the location they were sampled from, but it’s unclear whether other measurable features will be retained. For example, if an AMP with highly specific activity particular species or Gram classification is used as the reference, will the generated AMPs also selectively kill? The above-mentioned possibility of pairing with PepVAE3 with further compiled non-*E. coli* data for likely allowing for improved generation of AMPs targeting species of interest leads us to speculate whether this framework can result in a species-level granularity in terms of activity or whether other techniques would be better suited for this goal. In addition, we highlight that this study utilized a vanilla LSTM VAE that was used previously, not a more sophisticated variant, such as a conditional VAE (CVAE) for AMP generation (Das, Wadhawan et al. 2018). It’s unclear whether the results would be improved by conditioning the latent space on MIC information without an explicit comparison paired with experimental validation; in future studies we plan to investigate this, comparing the output of CVAE to PepVAE3, as well as other generative models to determine which performs the best for controlled AMP sequence generation.

## Methods

### Dataset and analysis of sequence characteristics

The dataset used in this study was based on Giant Repository of AMP Activity (GRAMPA) as described by Witten *et al.* (Witten and Witten 2019), with some modifications. Witten et al. scraped all data from APD (Wang, Li et al. 2016), DADP (Novković, Simunić et al. 2012), DBAASP (Pirtskhalava, Gabrielian et al. 2016), DRAMP (Fan, Sun et al. 2016), and YADAMP (Piotto, Sessa et al. 2012), each accessed in Spring 2018, resulting in a combined 6760 unique AMP sequences and 51345 MIC values, and is publicly available on GitHub: https://github.com/zswitten/Antimicrobial-Peptides. MIC values were independently spot-checked and confirmed; however, methods vary widely between publications, suggesting MIC values herein should be interpreted as approximations of activity. Since MICs determined against *E. coli* were the most-commonly available (**Figure S1A**), these were used for the study. To avoid issues with synthesis, the dataset was further modified by excluding peptides with cysteine and avoiding recorded modifications. For ease of synthesis and to keep costs low, sequences ≥ 40 amino acids in length (representing 3.1% of sequences) were excluded; see **Figure S1B-C** for the length distribution before and after this step). Since only the sequences and MIC values were needed, all other data from the modified GRAMPA dataset was removed. All MIC values were log μM transformed as done previously (Witten and Witten 2019). The sequences were tokenized, <end> token appended to each, and represented by a one-hot encoding scheme using binary vectors with length equal to the size of the amino acid vocabulary: the stopping token <end>, a, d, e, f, g, h, i, k, l, m, n, p, q, r, s, t, v, w, y, and a padding character. This resulted in a 3D data matrix of dimension 3280, 21, and 41 for the number of sequences, length of the vocabulary, and feature vector length, respectively. This process was repeated for *S. aureus* and *P. aeruginosa* identically to *E. coli.* The final 3D data matrices for *S. aureus* and *P. aeruginosa* had 2974 and 1968 sequences, respectively, with 21 and 41 length of the vocabulary and feature vector length. The *S. aureus* and *P. aeruginosa* datasets were only used for training of the MIC prediction regression models. Secondary structure of AMPs was predicted using the PredictHEC function from the DECIPHER R package within Bioconductor, providing the probability of helix, beta sheet, or coil (H, E, or C) (Wright 2016). PredictHEC makes use of the GOR IV algorithm (Garnier, Gibrat et al. 1996). This method is one of the best-performing that uses only the primary sequence and doesn’t require input of other sequences. Welch’s t-test via NumPy (Walt, Colbert et al. 2011) was used throughout for sample comparison with a significance threshold of 0.01 unless otherwise noted.

### Variational autoencoder

The architecture of the VAE was implemented as described by Bowman *et al.* (Bowman, Vilnis et al. 2015), as described by Dean and Walper (Dean and Walper 2020) with minor modifications. The loss function was comprised of reconstruction loss and KL loss to penalize poor reconstruction of the data by the decoder and encoder output representations of z (latent space variables) that are different from a standard normal distribution. The preprocessed data was encoded into vectors using a LSTM network. The encoder LSTM was paired with a decoder LSTM in order to do sequence-to-sequence learning. The decoder results were converted from binary one-hot encoded vectors back to peptide sequences. Training stoppage criteria was met when loss values did not decrease >0.0001 for five consecutive epochs. The VAE was trained using the Keras (Chollet 2015) library with a TensorFlow (Abadi, Barham et al. 2016) backend, and used the Adam optimizer. The number of neurons for the LSTM layers found in both encoder and decoder were both set to 1024. The number of latent dimensions was set to 50. All models were trained on an Ubuntu workstation with a Nvidia GeForce GTX1070 GPU. The LSTM network used in the decoder-encoder are stochastic – decoding from the same point in latent space may result in a different peptide being generated and is dependent on the random seed set prior to running. Models were saved to binary files and are available upon request.

### New sequence generation

Cosine similarity was used to compare latent space vectors generated by the VAE. To carry out cosine similarity calculations for each vector encoded, NumPy dot and norm functions were used which follow the notation:

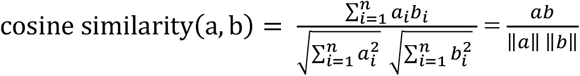

where a and b are vectors; ||*a*|| and ||*b*|| are Euclidean (L^2^) norms of vectors a=(a_1_, a_2_, …, a_n_) and b=(b_1_, b_2_, …, b_n_) (Han, Kamber et al. 2012), where each vector (b) was compared to the same reference (a): the vector representation of VLNENLLA, caseicin B-B1. Selected for its relative inactivity, caseicin B-B1 was reported to show a MIC of ≥ 1.25 mM against *E. coli* NCIMB 11843 in a study of caseicin B (Norberg, O’Connor et al. 2011).

Using the cosine similarity values, the five nearest to the caseicin B-B1 vector (Group A) and the five furthest from caseicin B-B1 (Group B) were identified. Around each of these vectors (*ν*), new vectors were sampled by selecting random points from a normal distribution. In order to accommodate the relative variation of the latent codes, we denote w_ij_ as a new vector with the following equation,

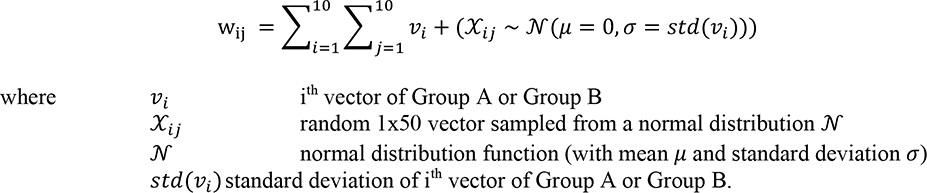

The *μ* was set to 0 and *σ* equal to the standard deviation of the vectors from Group A and Group B (*std*(*ν*)). The resulting random 1×50 vector 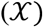 was then added to the input vector, resulting in a new vector (*w*). Using this method, 10 new vectors were sampled for Group A and Group B, and all vectors were decoded to new peptide sequences. Following removal of duplicate sequences and those already present in the modified GRAMPA dataset, 38 remained (sequences and other information available in **Table 1**).

### Latent space visualization

For dimensionality reduction PCA, t-SNE, UMAP – each with two components – was used. PCA and t-SNE used were imported from Scikit-learn, while UMAP was from McInnes *et al.* (McInnes, Healy et al. 2018). For t-SNE, perplexity set to 30, and learning rate set to 100. UMAP was performed using Bray-Curtis Similarity as the metric, with default settings. The MIC thresholds for coloring were: < 0.2 log μM is shown in blue, >2 log μM was set to red, and those with values ≥ 0.2 and ≤ 2 were set to light gray. To visualize the cosine similarity values of each encoded vector in latent space, the vector for each peptide was colored according to cosine similiartity value on a 2D t-SNE projection. To highlight the locations of Group A and Group B, black and white stars were placed at the points representing the peptides caseicin B-B1 and trialysin peptide P4, respectively.

### Regression models

Eight different regression models for predicting AMP MIC values: convolution neural network (CNN) as implemented by (Witten and Witten 2019), Elastic Net (ENet), Gradient Boosting (GB), Kernel Ridge (KR), Lasso, and Random Forest (RF) models used were from the Python Scikit-learn library (Pedregosa, Varoquaux et al. 2011), while Light Gradient Boosting Machine (LGBM) and EXtreme Gradient Boosting (XGB) used lightgbm (Ke, Meng et al. 2017) and xgboost (Chen and Guestrin 2016) libraries, respectively. The model parameters for each are provided in Supplemental Material. The data used for the regression models was the same as described in the *Dataset and analysis of sequence characteristics* section above, prior to one-hot encoding. The input peptides sequences were encoded numerically to vectors, each amino acid or padding characters – which were appended to the end vector below the maximum length (40) – receiving a unique number. The data was randomly shuffled and split into training:test sets at a 90:10 ratio. Initial comparison of the eight different regression models for predicting AMP MIC values against *E. coli* was performed by calculating RMSE and R^2^ values from actual MIC (log μM) vs. predicted on a holdout test dataset. From these the top performing four – GB, RF, LGBM, and XGB – were examined with shuffled split cross validation in each case (n=25) for predicting the MICs in the *E. coli, S. aureus,* and *P. aeruginosa* datasets. The top-performing MIC predictor for each organism was selected by lowest median RMSE.

### Minimum inhibitory concentration (MIC) assays

Minimum inhibitory concentration (MIC) measurements values were measured using broth dilution method for AMPs (Wiegand, Hilpert et al. 2008). Peptides synthesized for use in this study are listed in **Table 1**. Peptides were synthesized by Genscript, Inc. (Piscataway, NJ, USA) and each confirmed to have greater than 80% purity. Lyophilized peptides were solubilized in water, aliquoted, and stored at −20 °C. Overnight cultures of *E. coli* K-12, *Staphylococcus aureus* ATCC 12600, and *P. aeruginosa* 27853 were grown in Mueller Hinton II Broth, (BD, San Jose, CA, USA) at 37 °C. Cultures were diluted to a final concentration of approximately 5 × 10^5^ CFU/mL into fresh broth. An inoculum volume of 100 μl was added to each well of a 96-well non-treated polystyrene plate (Celltreat Scientific Products, Pepperell, MA, USA) and 10 μL of the peptide, which was diluted in series so that final peptide concentrations examined ranged from 128 μM to 0.5 μM. After incubation at 37 °C for 24 h, the MIC was determined by OD_600_ measurement using a BioTek Synergy Neo2 plate reader (Winooski, VT, USA) to identify the lowest concentration of peptide which inhibit growth. Statistical analysis of predicted and experimental MIC data was performed using the Fisher’s Exact Test from stats package in R (Team 2013).

### Circular dichroism

Circular dichroism (CD) spectra of the generated AMPs were obtained using a Jasco J-815 spectropolarimeter. Samples were allowed to equilibrate to 20 °C prior to data collection in a 0.1 cm path length cuvette, with a chamber temperature of 20 °C throughout each scan. Spectra were collected from 195 to 260 nm in 0.1-nm intervals. Data shown is an average of three scans, performed for each sample. All AMPs were analyzed at 25 μM concentration in either 10 mM sodium phosphate (pH 7) or 60 mM sodium dodecyl sulfate (SDS) in 10 mM sodium phosphate, both from Sigma (St. Louis, Missouri, USA). Baselines obtained prior to the experiment with peptide-free buffers were subtracted from each scan. Mean residue molar ellipticity (MRME) was calculated as follows:

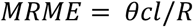

Where *θ* is ellipticity (mdeg), *c* is peptide concentration (mol/L), *l* is cell path length (cm), and *R* is the length of the peptide. MRME is presented multiplied by 1000 to improve clarity.

## Acknowledgements

We acknowledge funding support through the Jerome and Isabella Karle Distinguished Scholar Fellowship provided by the Naval Research Laboratory, base funds of the Naval Research Laboratory (WU# 1V33), and funds from the Defense Threat Reduction Agency (HDTRA1033536).

## Supplemental Material

### Methods

#### Peptide characterization

The characteristics of sequences in the starting dataset and generated AMPs, including peptide length, amino acid composition, net charge, hydrophobicity, and hydrophobic moment, were assessed using the Python library modlAMP (Müller, Gabernet et al. 2017).

#### Preliminary methods comparisons

Machine learning methods were implemented using the MATLAB Statistics and Machine Learning Toolbox, and the Regression Learner app. The models tested included: Rational Quadratic Gaussian Process Regression, Fine Gaussian Support Vector Machine, Ensembles of Trees – Bagged, and Regression Trees – Coarse, as well as others part of the app. The Regression Learner app was used to select the most promising methods for further study. The use of structured data was necessary due to the default behavior of MATLAB that treats an array of character (char) data as a single string. The amino acid identities as the standard single-letter abbreviation were held in a table consisting of unit cell arrays which each contained a single char datum. To this was appended the response variable, the logarithm of the Minimum Inhibitory Concentration, log(MIC).

The amino acid sequences were of variable lengths. Since machine learning methods typically require equal length data, it is necessary to pad the data, either at the C-terminus, or by a multiple alignment method. C-terminal padding simply involves the addition of a non-coding character to the end of shorter peptide sequences to bring them all up to the length of the longest example. Multiple alignment was performed with the MATLAB function multialign from the Bioinformatics Toolbox. MATLAB machine learning treats the amino acid identities (as well as noncoding characters) as categorical type data and gives no special meaning to the actual amino acid.

#### Latent space analysis

As previously used to evaluate VAE-arrived clustering, methods described by Lopez *et al.* were used, with modification (Lopez, Regier et al. 2018). The 2D projections resulting from the dimensionality reduction methods PCA, t-SNE, and UMAP from the VAE-produced latent representation was used as input to the *K*-means algorithm and measure the overlap between the resulting clustering annotations and the pre-specified subpopulations (the < 0.2 log μM and >2 log μM labels) using Adjusted Rand index and Adjusted Mutual Information (AMI) measurements as metrics.

#### Minimum inhibitory concentration (MIC) assay analysis

Statistical analysis of predicted and experimental MIC data was performed using receiver operating characteristic (ROC) curve analysis and area under the curve (AUC) calculation from the pROC package (Robin, Turck et al. 2011) following categorization of predicted and experimental MICs into either >128 μM or ≤ 128 μM.

#### Circular dichroism analysis

Estimates of secondary structure from circular dichroism data. Measured ellipticity (mdeg) converted to Δ*ε* was input into the circular dichroism data analysis program Beta Structure Selection (http://bestsel.elte.hu) from which helix, antiparallel, parallel, turn, and other were obtained (Micsonai, Wien et al. 2018). Following summation of antiparallel, parallel, and turn into Sheet, the results were plotted for both groups A and B.

---

**Table S1.**
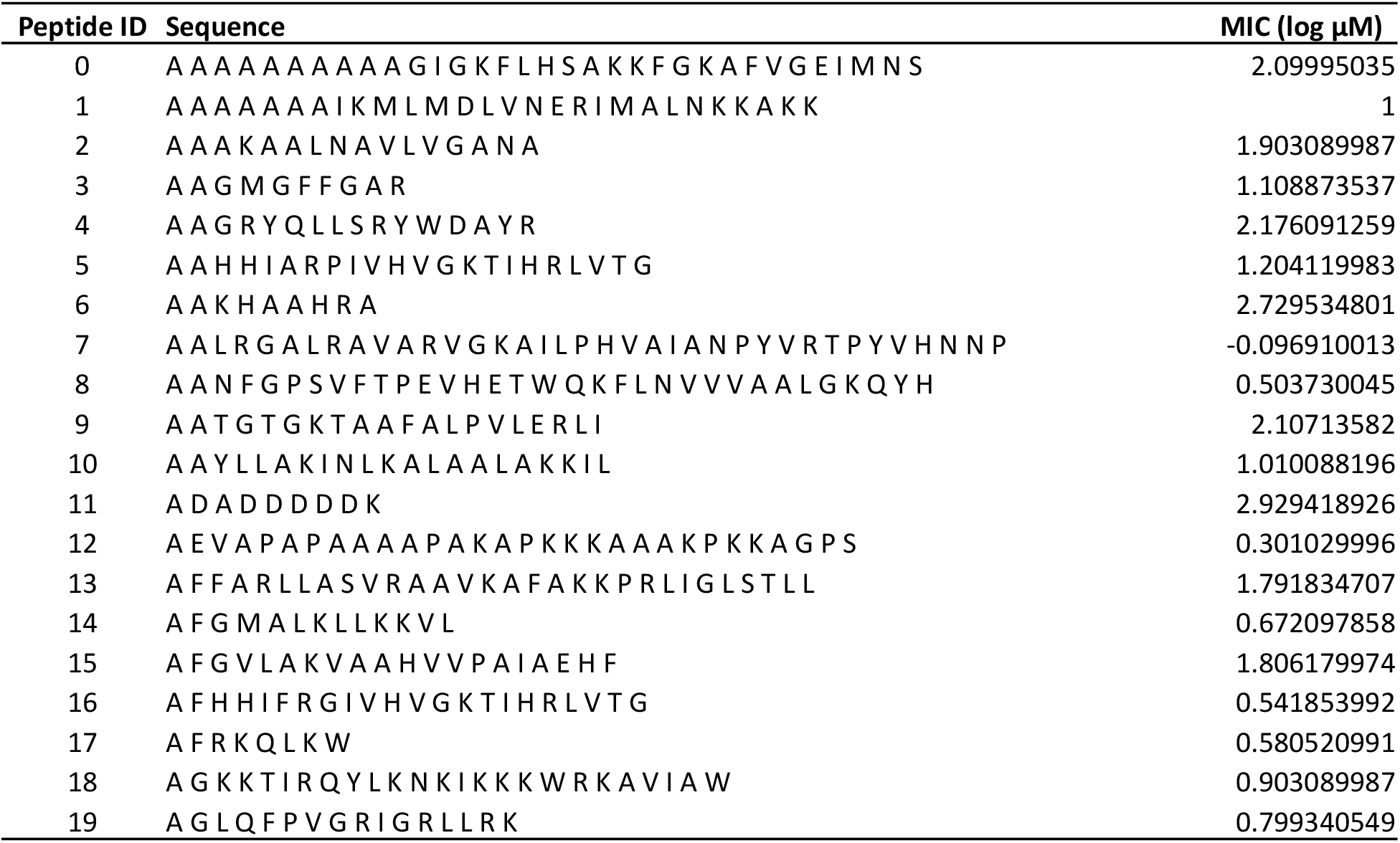
*E. coli* AMP and MIC values dataset used for training the VAE.

**Table S2.**
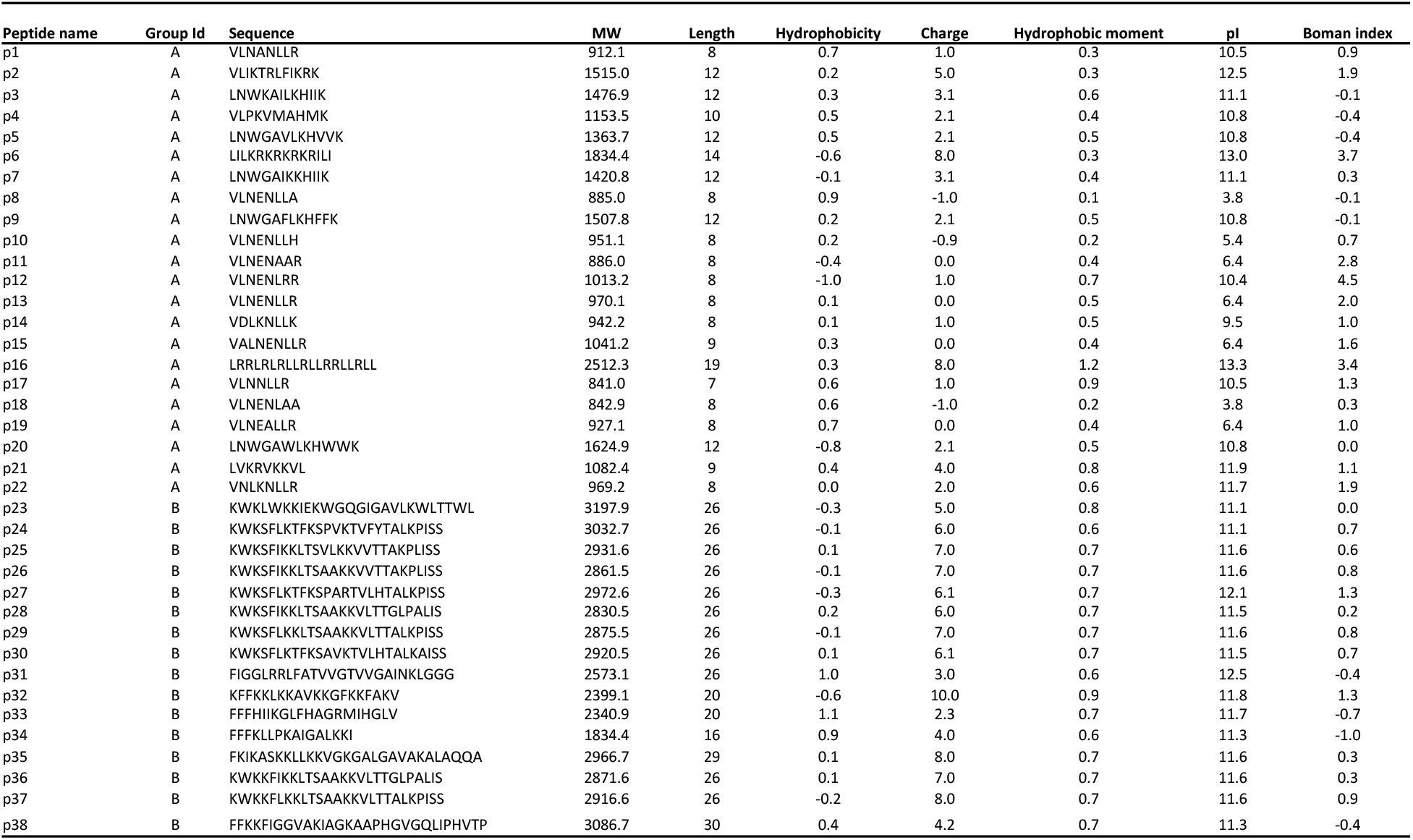
Characteristics of the peptides in the study.

**Figure S1.**
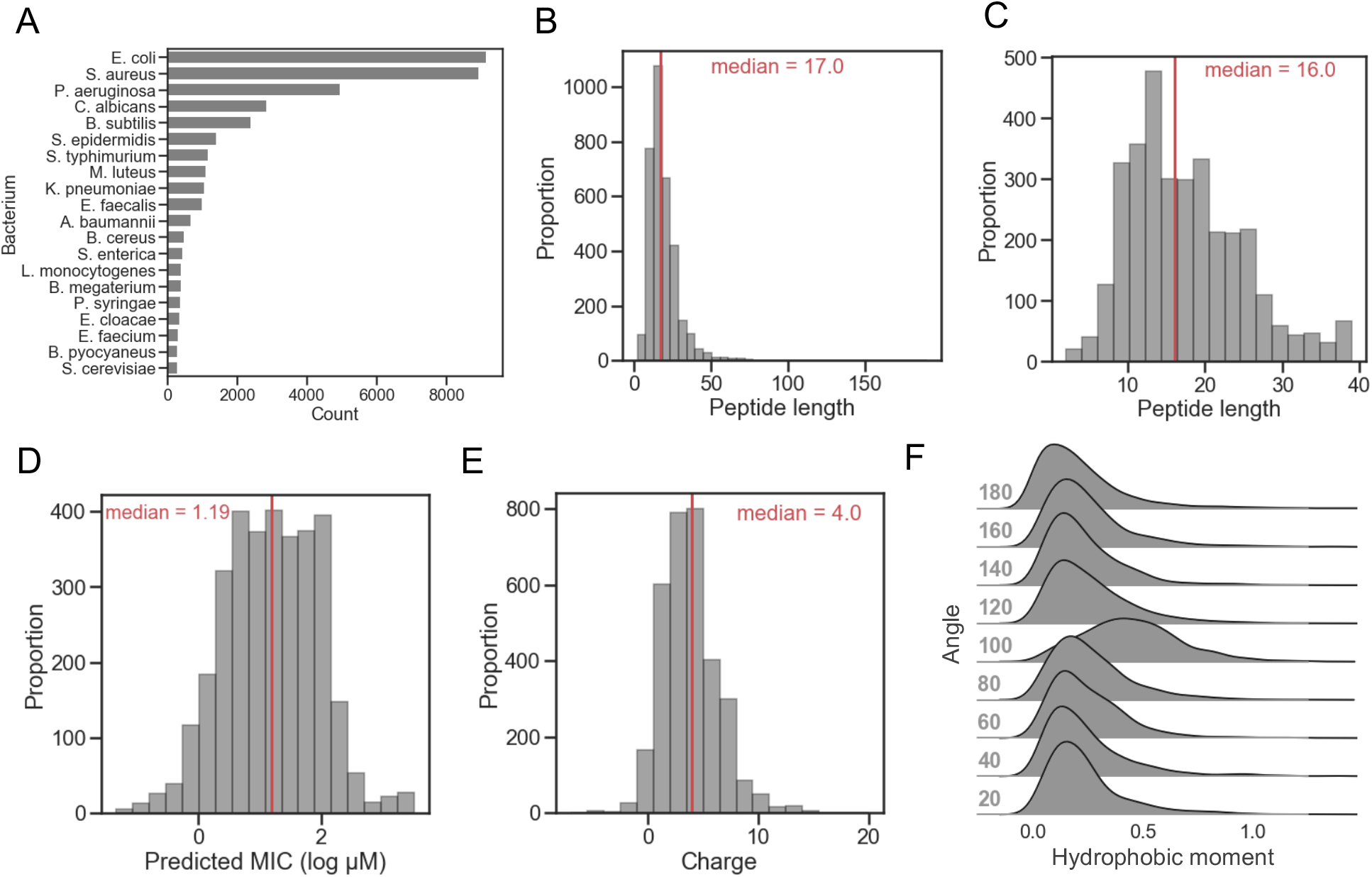
Characteristics of the AMP dataset. A) Top 20 bacteria tested against ordered by count in the initial GRAMPA dataset. *E. coli*, *S. aureus,* and *P. aeruginosa* are most commonly tested, with *E. coli* the most counted at 9150. B) Peptide length distribution of AMPs (without cysteine) tested against *E. coli*. C) Peptide length distribution of AMPs (without cysteine) tested against *E. coli*, with AMPs of length ≥ 40 removed. D, E, and F show the distribution of MIC (log μM), charge, and hydrophobic moments at different angles, respectively, of the filtered peptides represented in panel C. Medians for distributions shown in B, C, D, and E are indicated with a red line.

**Figure S2.**
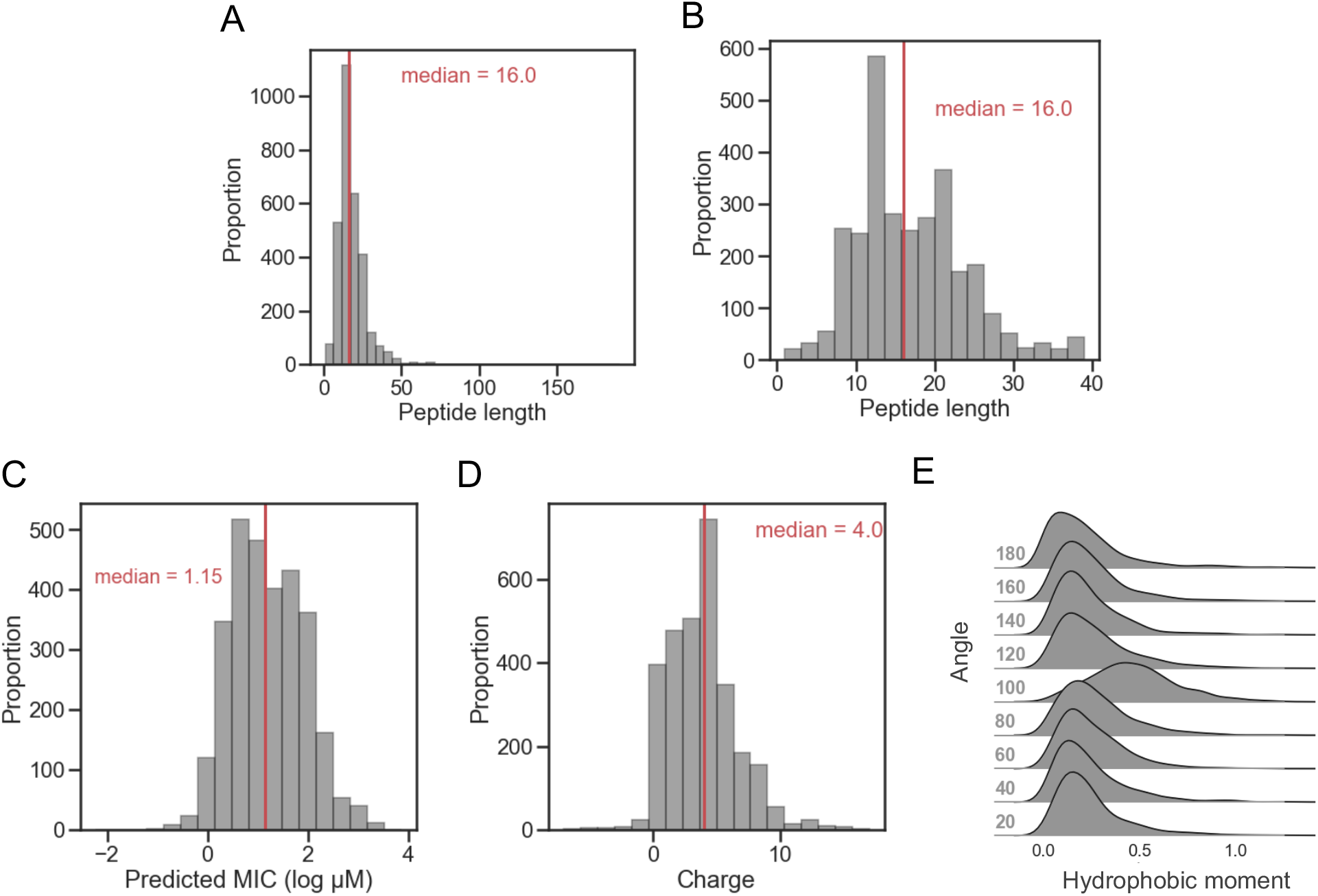
Characteristics of the AMP dataset for *S. aureus*. A) Peptide length distribution of AMPs (without cysteine) tested against *S. aureus*. B) Peptide length distribution of AMPs (without cysteine) tested against *S. aureus*, with AMPs of length ≥ 40 removed. C, D, and E show the distribution of MIC (log μM), charge, and hydrophobic moments at different angles, respectively, of the filtered peptides represented in panel C. Medians for distributions shown in A, B, C, and D are indicated with a red line.

**Figure S3.**
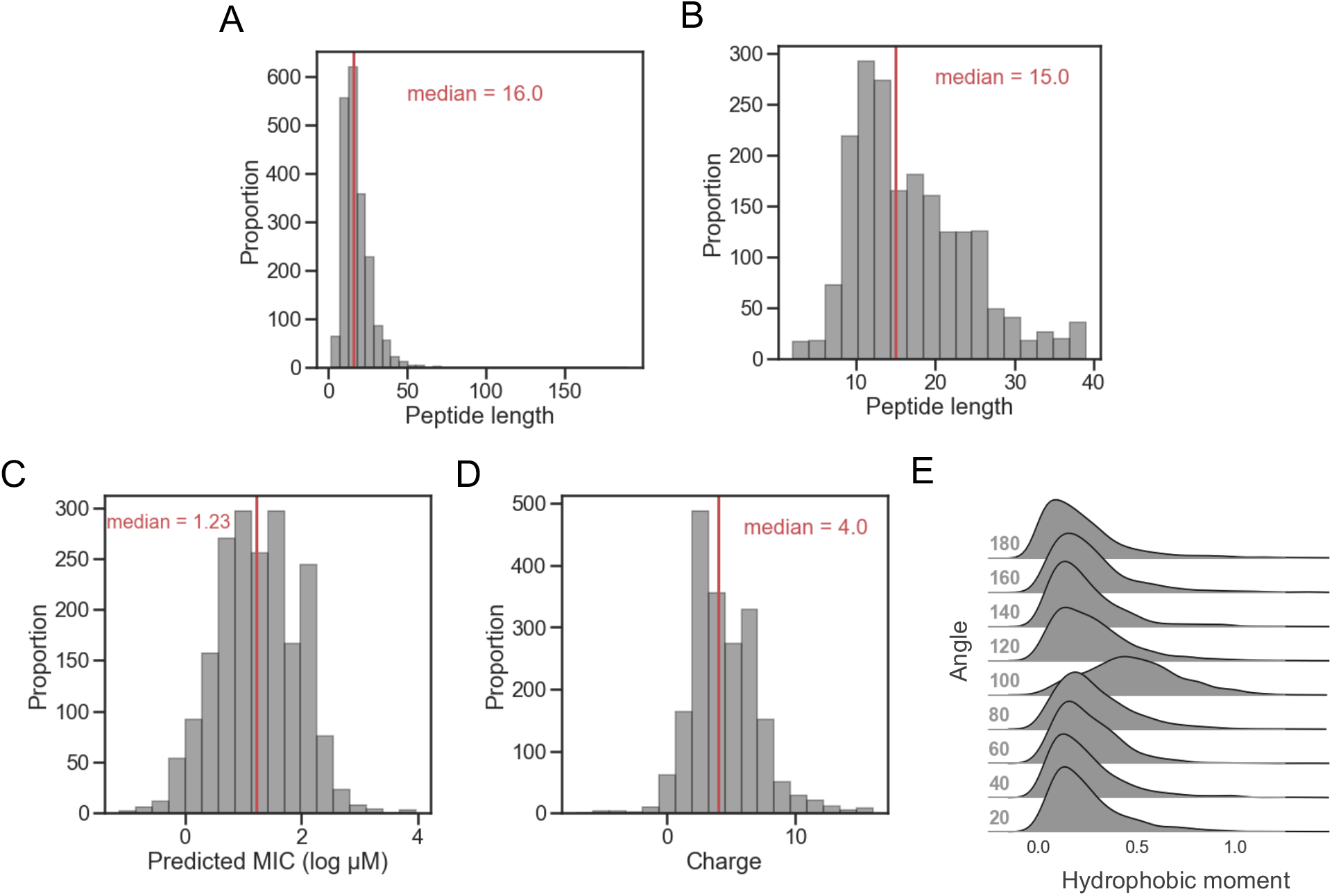
Characteristics of the AMP dataset for *P. aeruginosa*. A) Peptide length distribution of AMPs (without cysteine) tested against *P. aeruginosa*. B) Peptide length distribution of AMPs (without cysteine) tested against *P. aeruginosa*, with AMPs of length ≥ 40 removed. C, D, and E show the distribution of MIC (log μM), charge, and hydrophobic moments at different angles, respectively, of the filtered peptides represented in panel C. Medians for distributions shown in A, B, C, and D are indicated with a red line.

**Figure S4.**
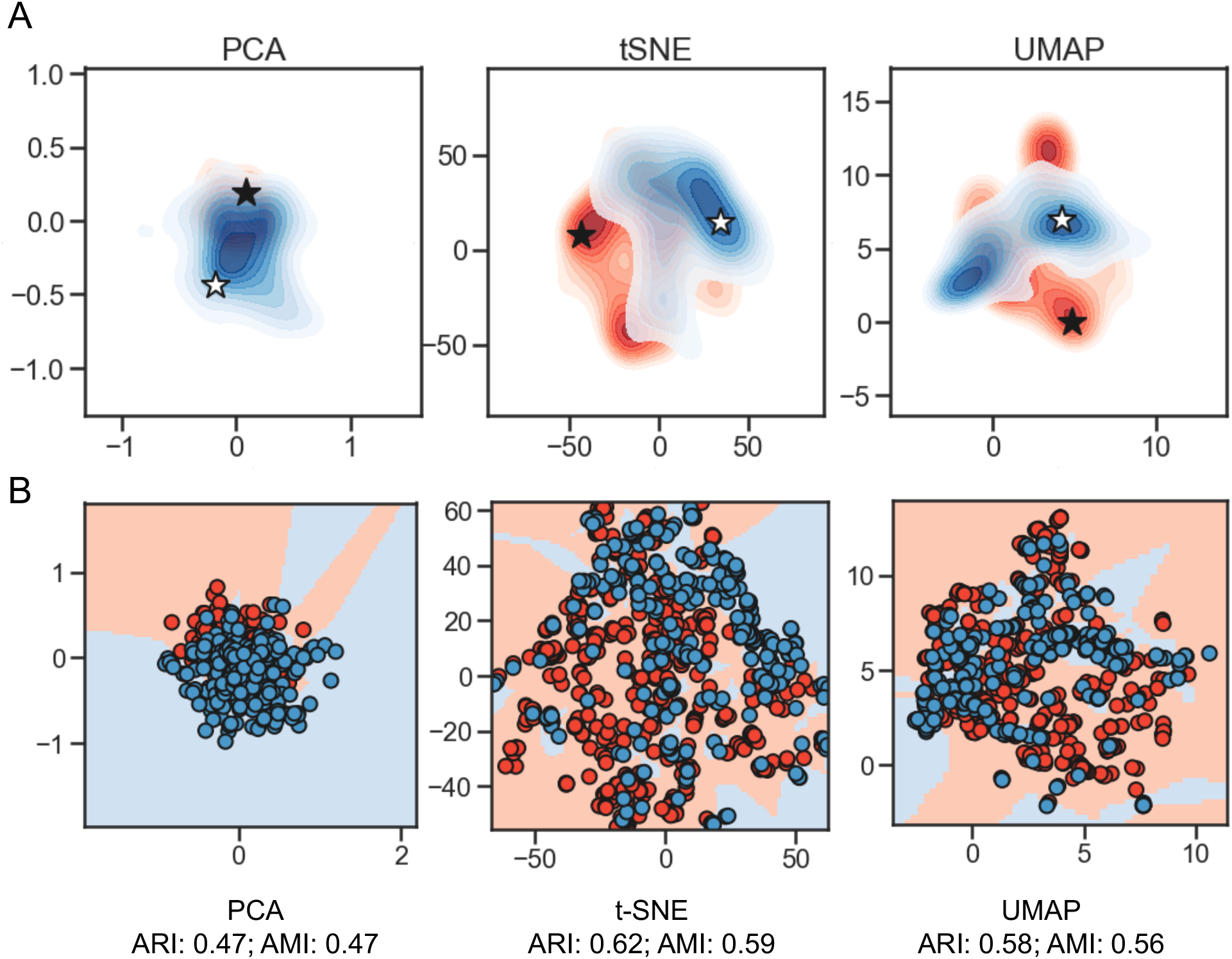
Dimensionality reduction for visualization of the 50-dimension latent space. A) Three different techniques were used: PCA, t-SNE, and UMAP (each with two components). PCA and t-SNE were imported from Scikit-learn; UMAP was from McInnes *et al.* (McInnes, Healy et al. 2018). The MIC thresholds for coloring were: < 0.2 log μM is shown in blue, >2 log μM was set to red, and those with values ≥ 0.2 and ≤ 2 were set to light gray. For t-SNE, perplexity set to 30, and learning rate set to 100. UMAP was performed using Bray-Curtis Similarity as the metric, with default settings. The black star is located at the peptide VLNENLLA, and a white star is located at the embedding most distant in cosine similarity (encoding for the peptide KFGKIVGKVLKQLKKVSAVAKVAMKKG). B) Adjusted Rand index (ARI) and Adjusted Mutual Information (AMI) measurements (for both, higher is better). KNN predictions performed on the 2D projections resulting from the dimensionality reduction methods PCA, t-SNE, and UMAP shown in (A), compared with the true labels were used to calculate the Rand index values shown above each plot. Scatterplot points represent locations of encoded peptides, background coloring indicates location of the KNN decision boundary. MIC thresholds for coloring: < 0.2 log μM is shown in blue, >2 log μM in red.

**Figure S5.**
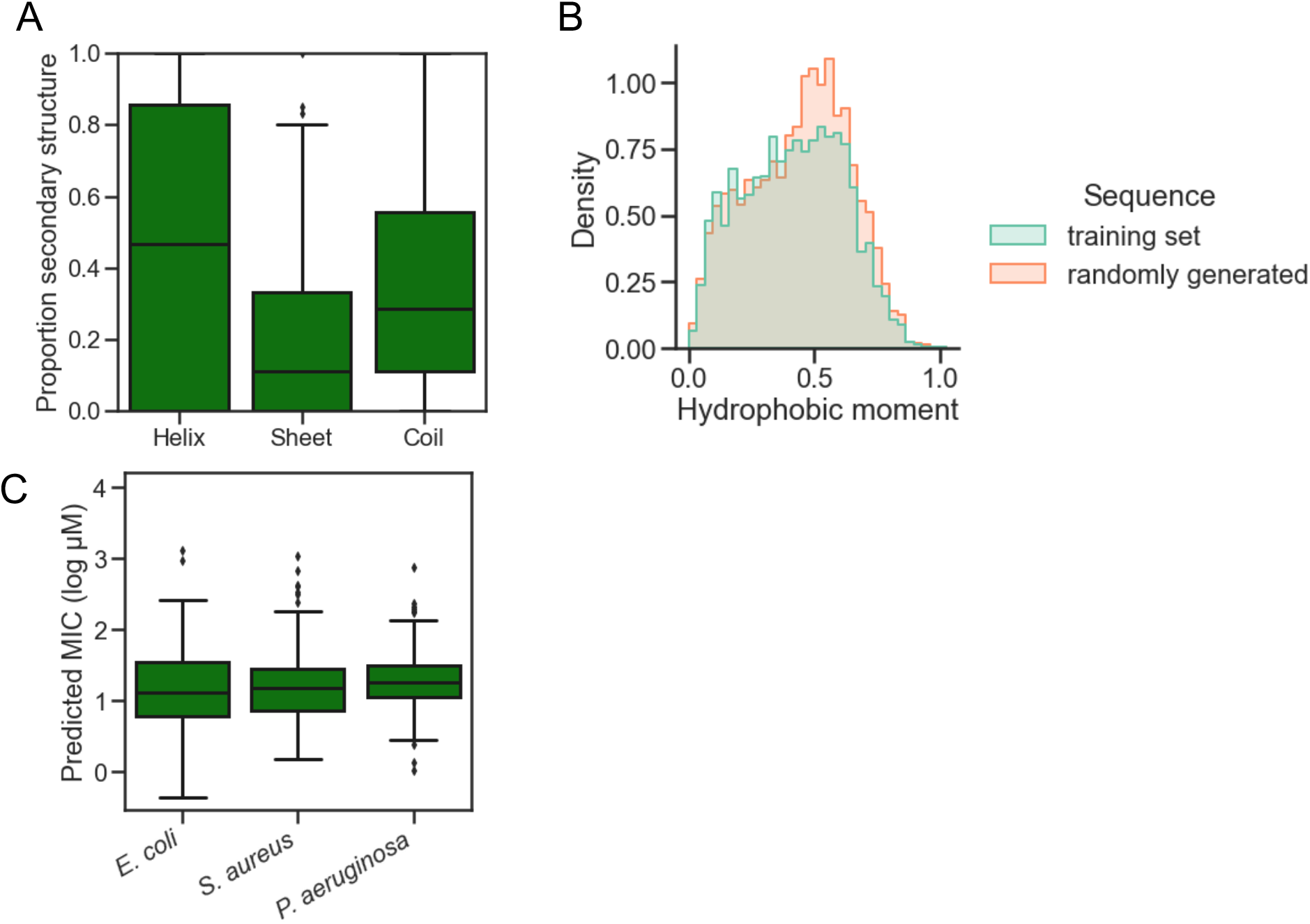
Analysis of randomly generated sequences. Points from latent space were selected at random and decoded to sequences. A) Boxplot of the predicted helix, sheet, and coil percentages calculated via the GOR IV algorithm. B) Density plot of the calculated hydrophobic moments obtained by applying the modlAMP function calculate_moment on the sequences from the randomly generated set and training set. C) A) Predicted MICs for the randomly generated set using the top-performing models for *E. coli*, *S. aureus*, and *P. aeruginosa*.

**Figure S6.**
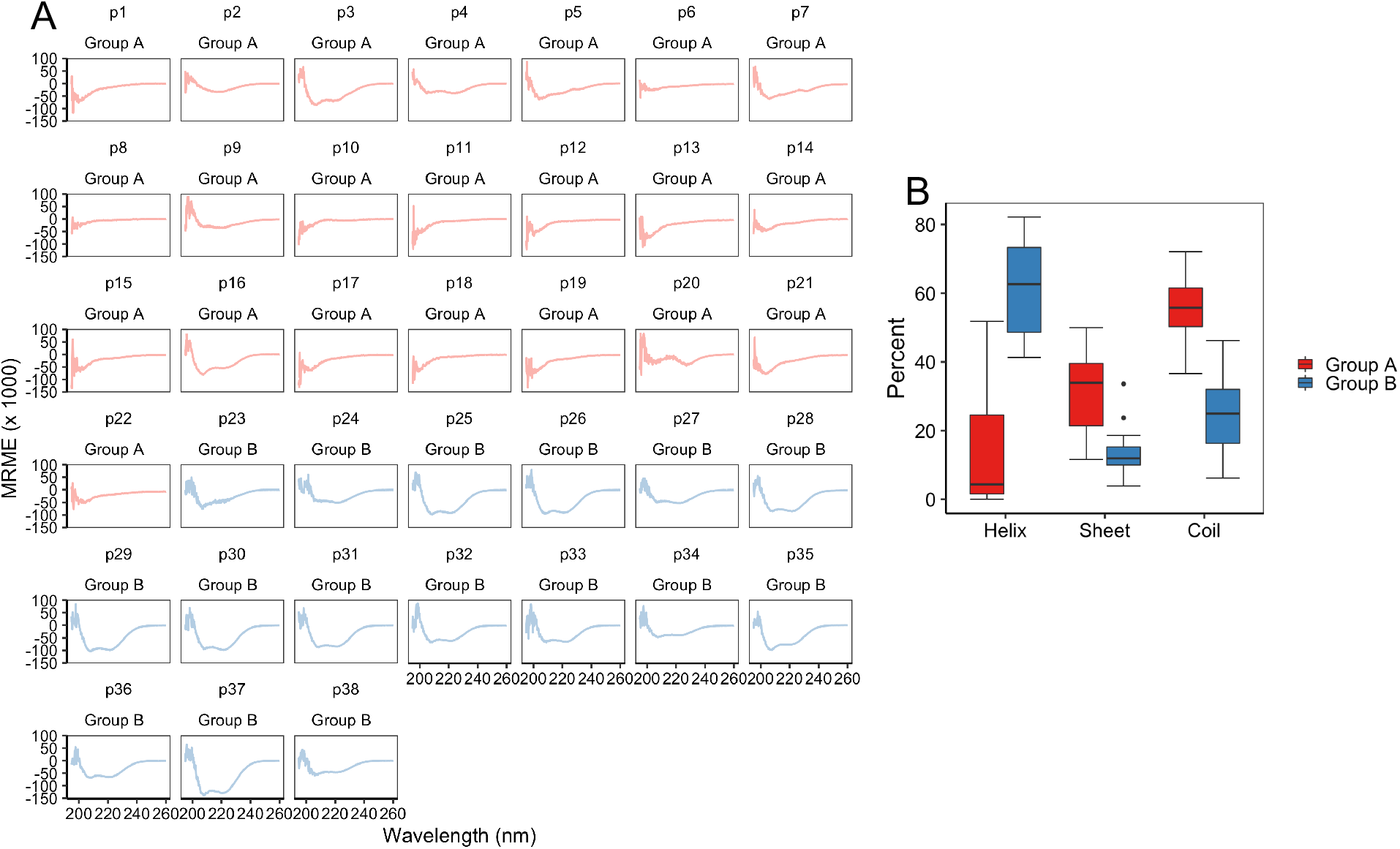
Circular dichroism scans and summary. A) Mean residue molar ellipticity (MRME) plots of peptides from group A and group B in the presence of membrane mimic 60 mM SDS. Each scan was averaged from three scans for each peptide with peptide-free buffer baseline scan subtracted. B) Estimates of secondary structure from circular dichroism data. The Beta Structure Selection CD analysis program was used to obtain helix, antiparallel, parallel, turn, and other. Following summation of antiparallel, parallel, and turn into Sheet, the results were plotted for both groups A and B.

**Figure S7.**
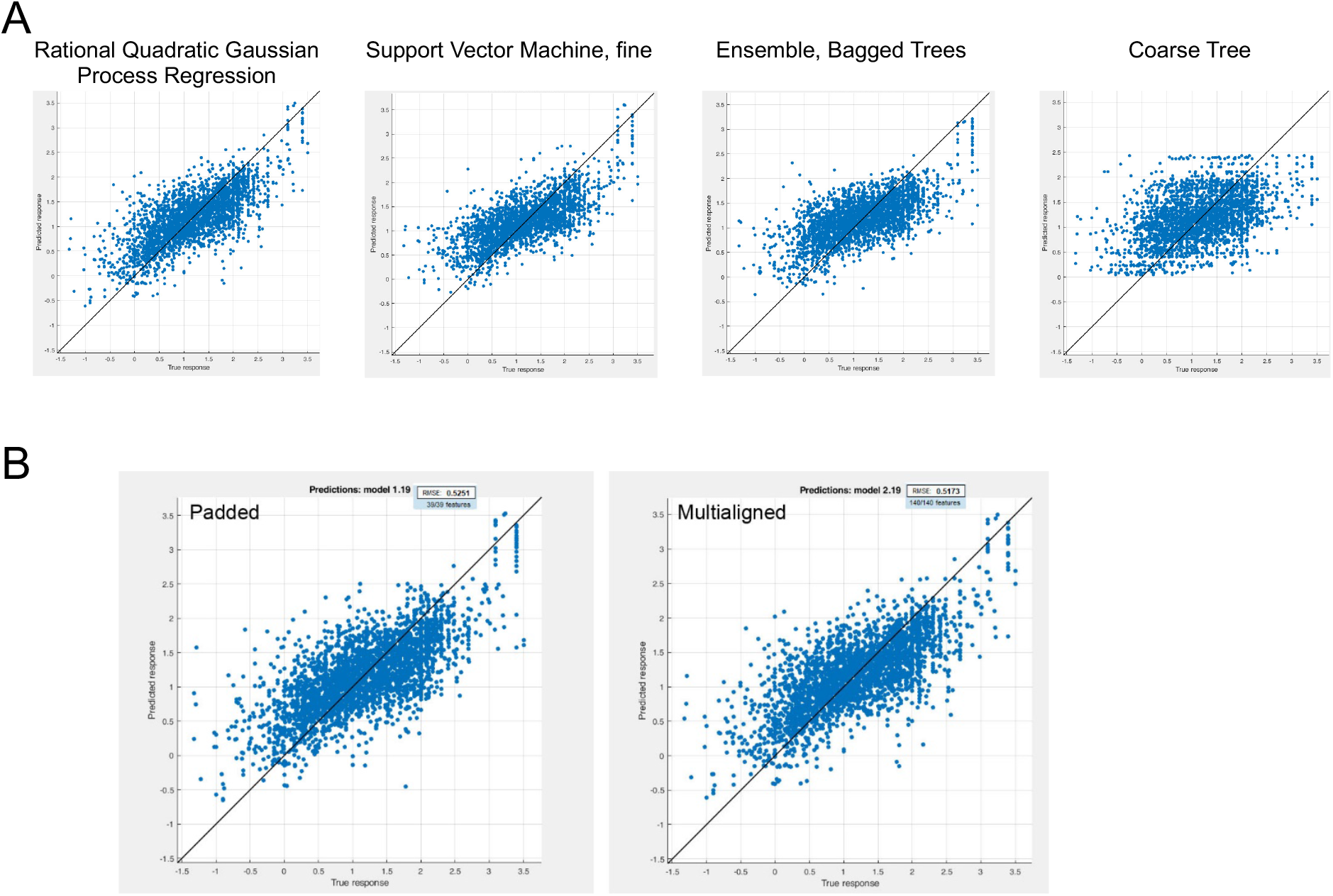
Regression model investigation with MATLAB. A) Representative performance of models trained on *E. coli* dataset from Statistics and Machine Learning Toolbox and Regression Learner app in MATLAB. RMSE values were: Rational Quadratic Gaussian Process Regression: 0.517, Fine Gaussian Support Vector Machine: 0.546, Ensembles of Trees – Bagged: 0.579, and Regression Trees – Coarse: 0.689. B) Comparison of Rational Quadratic Gaussian Process Regression performance using multi-aligned or padded sequences as input.

**Figure S8.**
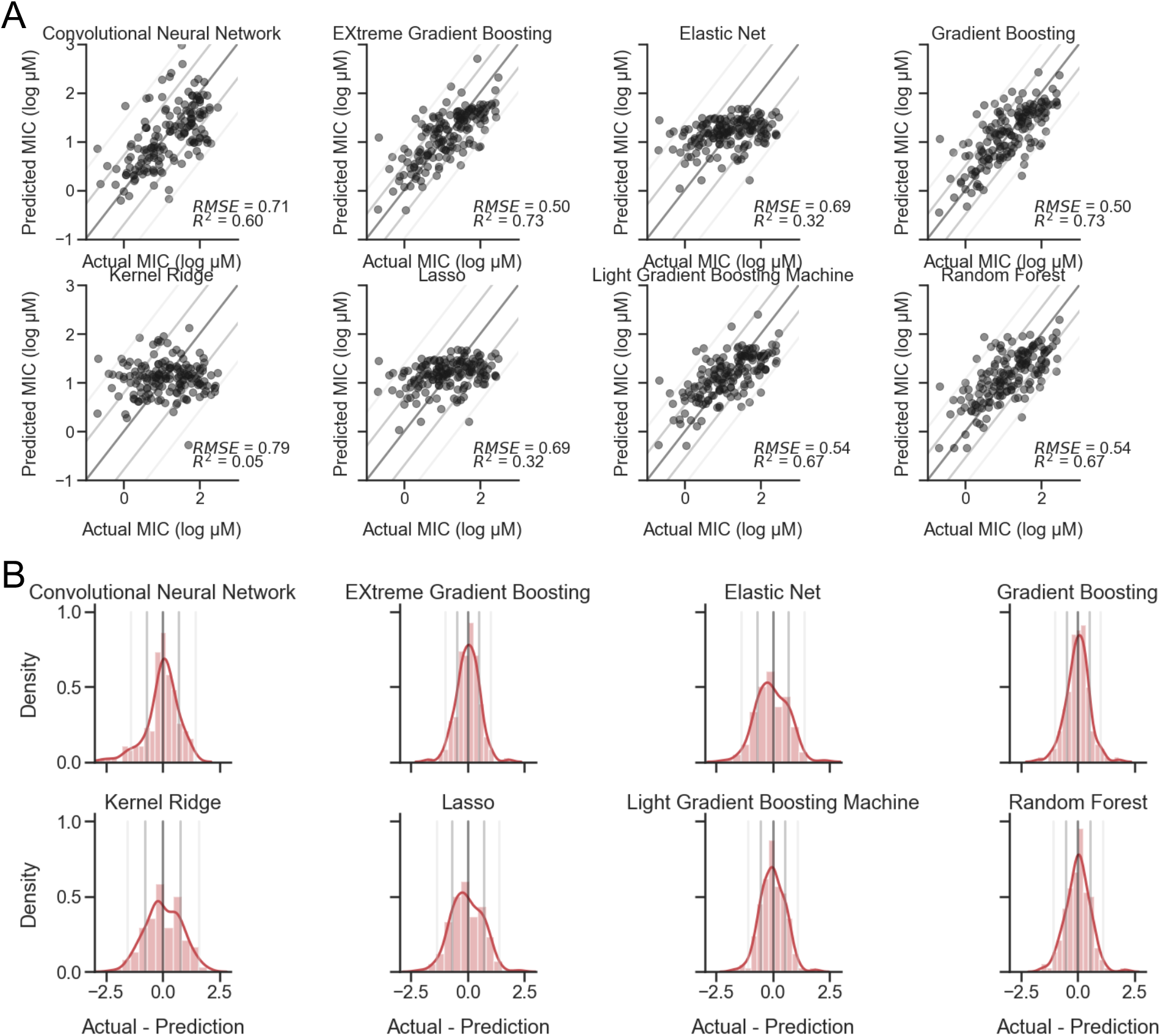
Initial regression model comparison. Eight different regression models for predicting AMP MIC values against *E. coli*. A) Representative scatterplots of Predicted vs. Actual MIC (log μM) of Convolutional Neural Network, Elastic Net, Gradient Boosting, Kernel Ridge, Lasso, and Random Forest, Light Gradient Boosting Machine, and EXtreme Gradient Boosting predictions on holdout test dataset with RMSE and R^2^ values displayed. Lines represent standard diagonals, in addition to the diagonal +/− 1 and 2 standard deviations of the points. B) Histograms with overlayed density curves of the Actual - Prediction difference of each model. Lines represent standard diagonals, in addition to the vertical midline +/− 1 and 2 standard deviations of the points.

**Figure S9.**
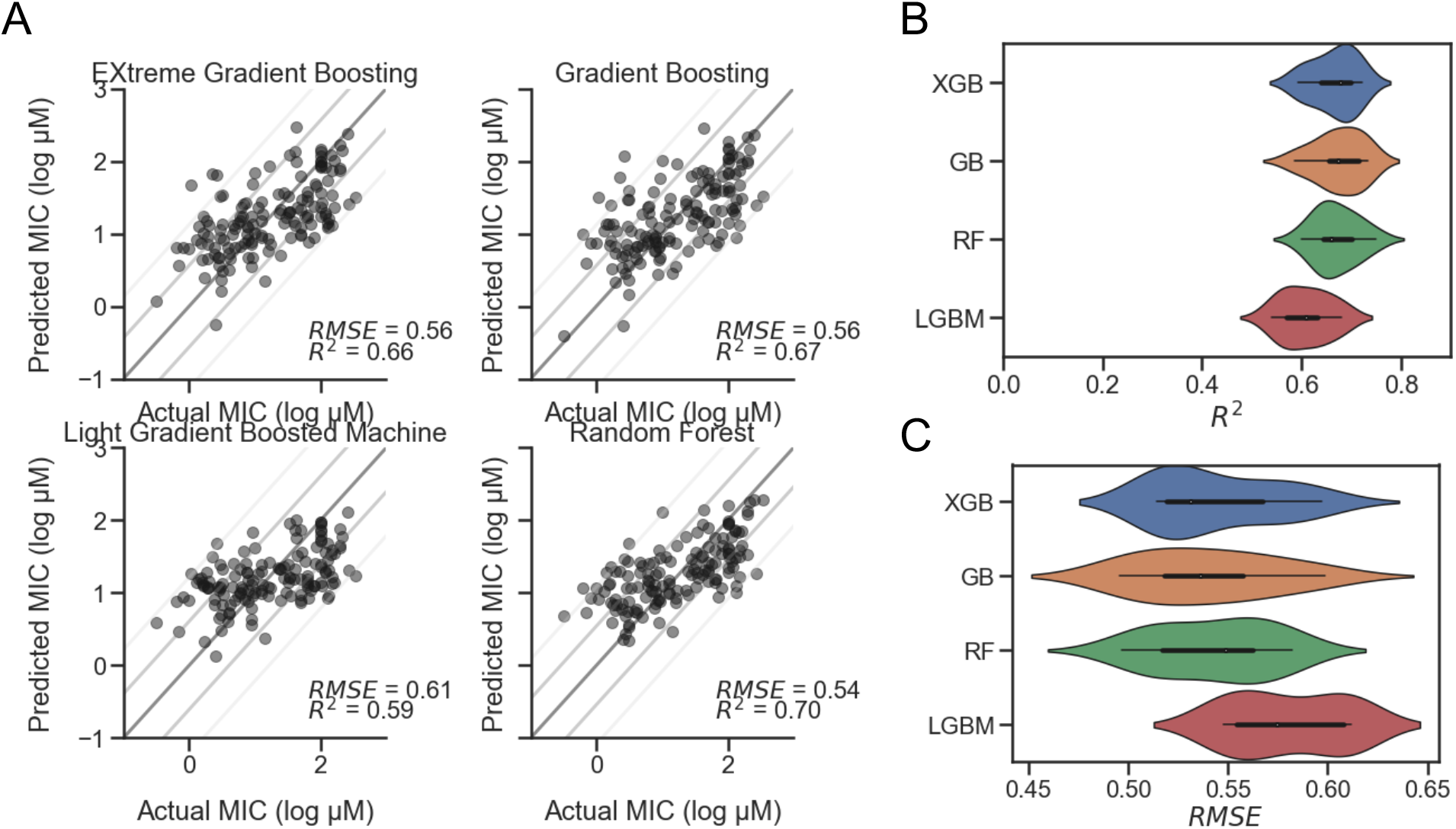
Comparison of MIC prediction models against *S. aureus*. Four regression models for predicting AMP MIC values against *S. aureus*. A) Representative scatterplots of Predicted vs. Actual MIC (log μM) of EXtreme Gradient Boosting (XGB), Gradient Boosting (GB), Light Gradient Boosting Machine (LGBM), and Random Forest (RF) predictions on holdout test dataset with RMSE and R^2^ values displayed. Lines represent standard diagonals, in addition to the diagonal +/− 1 and 2 standard deviations of the points. B) Results of cross validation using shuffled split (n=25) shown as a violin plot for each model, sorted from highest to lowest mean R^2^ value, with XGB at the top with a median of 0.7. C) Cross validation results for RMSE sorted from lowest to highest mean RMSE value, with XGB at the top with a median of 0.53.

**Figure S10.**
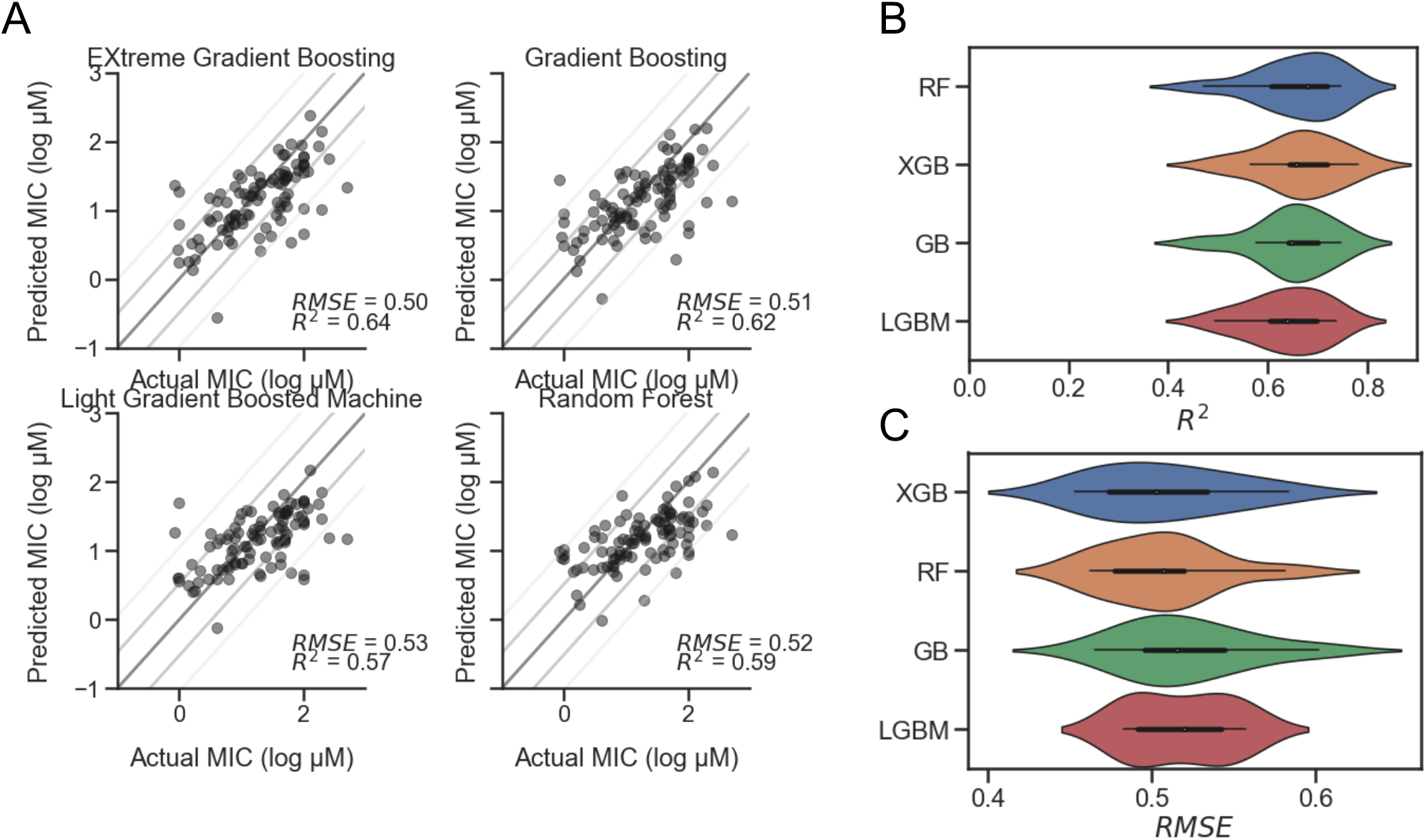
Comparison of MIC prediction models against *P. aeruginosa*. Four regression models for predicting AMP MIC values against *P. aeruginosa*. A) Representative scatterplots of Predicted vs. Actual MIC (log μM) of EXtreme Gradient Boosting (XGB), Gradient Boosting (GB), Light Gradient Boosting Machine (LGBM), and Random Forest (RF) predictions on holdout test dataset with RMSE and R^2^ values displayed. Lines represent standard diagonals, in addition to the diagonal +/− 1 and 2 standard deviations of the points. B) Results of cross validation using shuffled split (n=25) shown as a violin plot for each model, sorted from highest to lowest mean R^2^ value, with RF at the top with a median of 0.69. C) Cross validation results for RMSE sorted from lowest to highest mean RMSE value, with XGB at the top with a median of 0.5.

**Figure S11.**
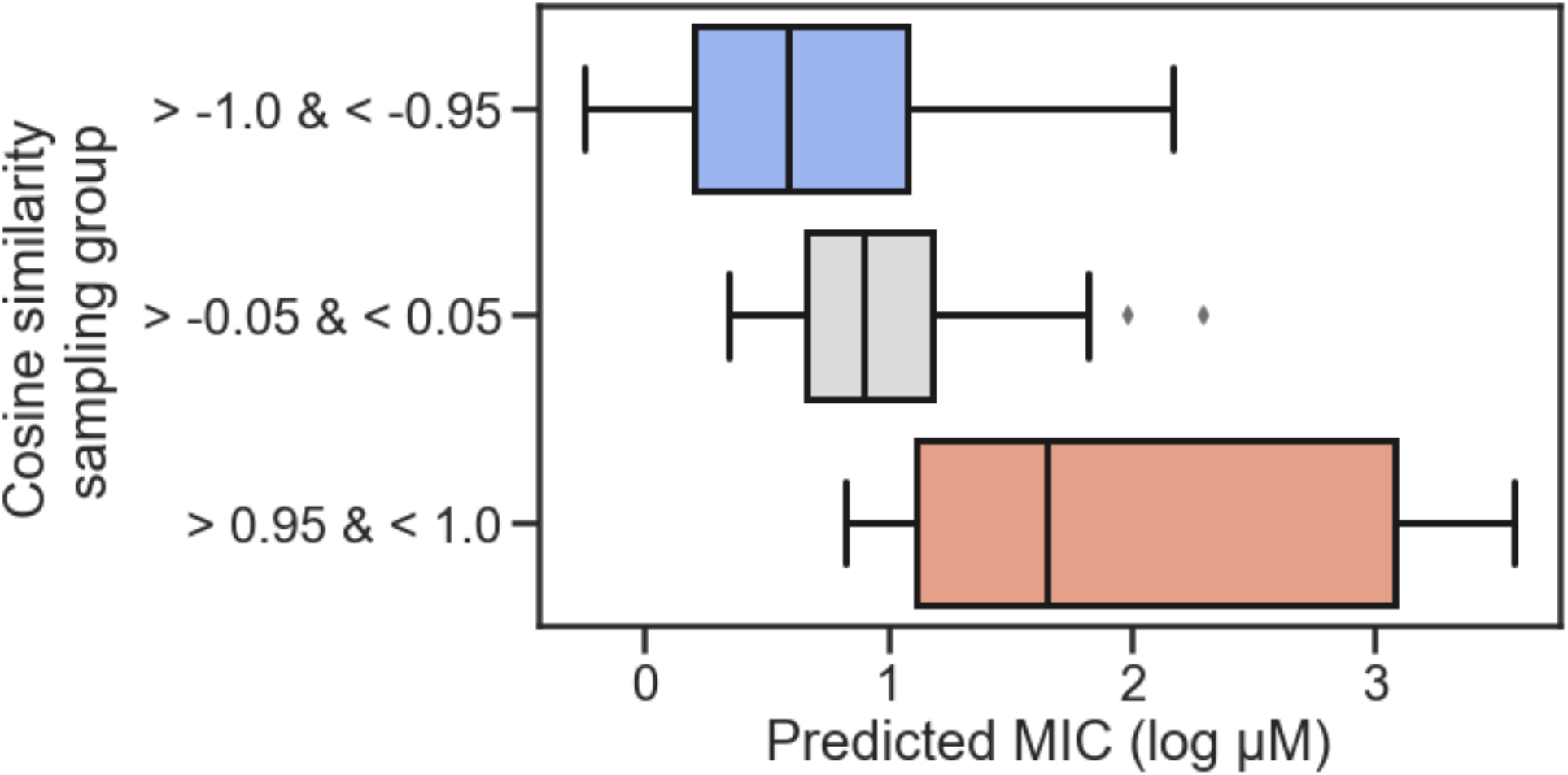
MIC predictions on cosine similarity sampling groups. Using the latent vector for VLNENLLA as a reference, vectors for each peptide were sorted by cosine similarity from −1.0 to 1.0 (1.0 being identical to the reference). Peptides were selected at random (n=100) from each of the following groups, filtering on cosine similarity: between −1.0 and −0.95 (> −1.0 & < −0.95), between −0.05 and 0.05 (> −0.05 & < 0.05), and between 0.95 and 1.0 (> 0.95 & < 1.0). New vectors (n=100) were generated near to the selected peptides and decoded to new sequences, to which the *E. coli* MIC prediction model was applied following removal of duplicate sequences. Boxplot summarizes the predicted MICs as a function of cosine similarity group.

**Figure S12.**
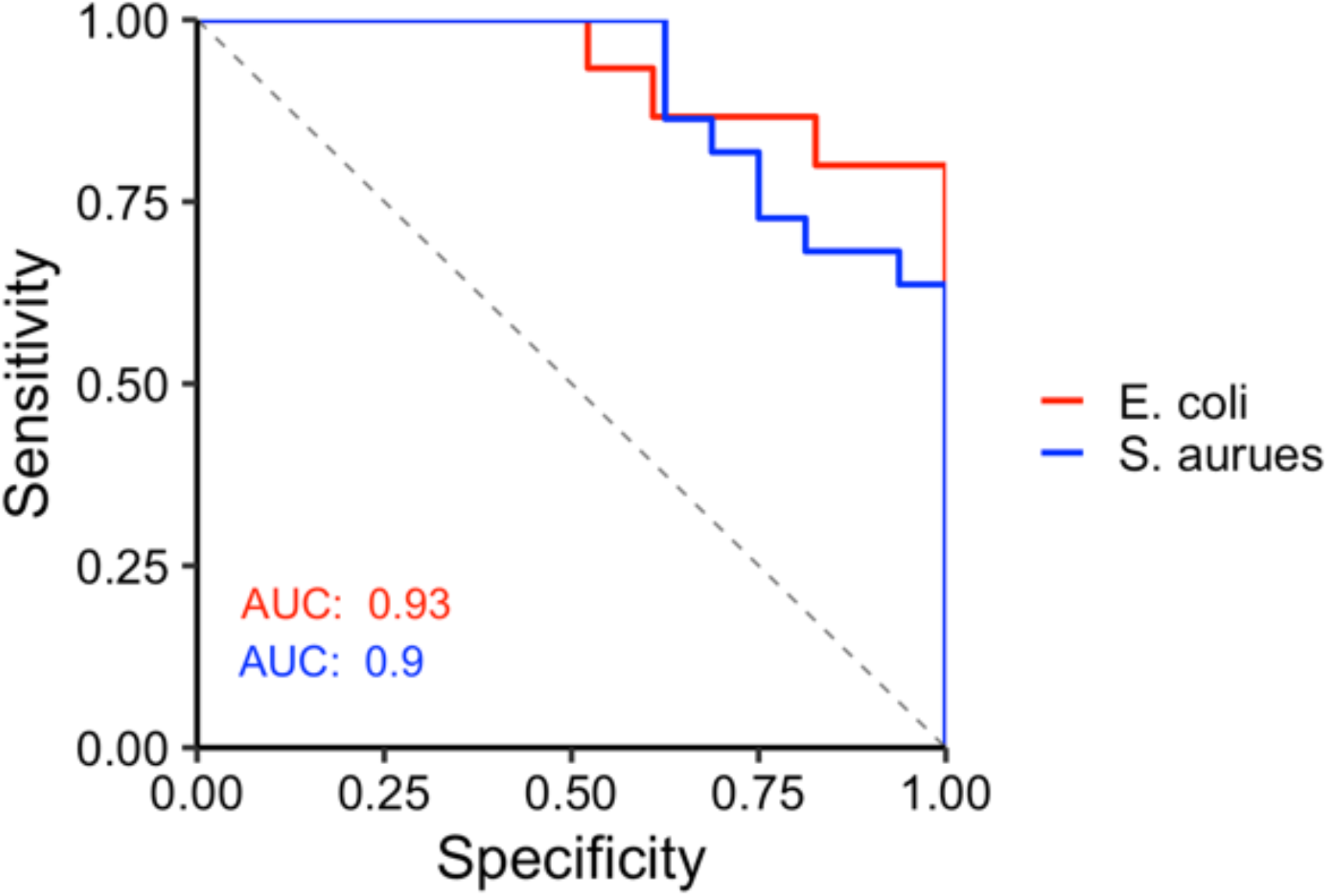
Receiver operating characteristic curve analysis of predicted and experimental MIC results. Receiver operating characteristic (ROC) analysis of predicted and experimental MICs of the 38 synthesized AMPs against *E. coli* and *S. aureus* when categorized into >128 and ≤128 μM yielded area under the curve (AUC) values of 0.93 and 0.9, respectively.

## Notes

### Competing Interest Statement

The authors have declared no competing interest.

